# Structural basis of the bacterial flagellar motor rotational switching

**DOI:** 10.1101/2024.04.30.591856

**Authors:** Jiaxing Tan, Ling Zhang, Xingtong Zhou, Siyu Han, Yan Zhou, Yongqun Zhu

## Abstract

The bacterial flagellar motor is a huge bidirectional rotary nanomachine that drives rotation of the flagellum for bacterial motility^1–5^. The cytoplasmic C ring of the flagellar motor functions as the switch complex for the rotational direction switching from counterclockwise to clockwise^2,3,6–9^. However, the structural basis of the rotational switching and how the C ring is assembled have long remained elusive. Here, we present two high-resolution cryo-electron microscopy structures of the C ring-containing flagellar motor-hook complex from *Salmonella* Typhimurium, which are in the default counterclockwise state and in a constitutively active CheY mutant-induced clockwise state, respectively. In both complexes, the C ring consists of four subrings, but is in two different conformations. The CheY proteins are bound into an open groove between two adjacent protomers on the surface of the middle subring of the C ring and interact with the FliG and FliM subunits. The binding of the CheY protein induces a significant upward shift of the C ring towards the MS ring and inward movements of its protomers towards the motor center, which eventually remodels the structures of the FliG subunits and reverses the orientations and surface electrostatic potential of the α_torque_ helices to trigger the counterclockwise-to-clockwise rotational switching. The conformational changes of the FliG subunits reveal that the stator units on the flagellar motors require a relocation process in the bacterial inner membrane during the rotational switching. This study provides unprecedented molecular insights into the rotational switching mechanism and a detailed overall structural view of the bacterial flagellar motors.

## Introduction

The bacterial flagellum is a proteinaceous organelle responsible for motility, which is crucial for bacterial infection, biofilm formation and survival in various environments^10–12^. The structure of the flagellum is made up of three distinct parts: the motor, the hook and the filament^4,13^. The motor is a bidirectional rotary nanomachine that provides the power for the rotation of the flagellum^3^. The hook functions as a joint that connects the motor to the filament, which acts as a propeller to propel bacteria for swimming in liquid medium and swarming on solid surface^3^. As the basal body of the flagellum, the motor spans both the inner and outer membranes, and is made up of the LP ring, MS ring, C ring, rod, export apparatus and several stator units^14–17^. The LP ring acts as a bushing to stabilize the rotation of the rod, which serves as the drive shaft. The MS ring is located on the inner membrane, houses the export apparatus and forms the assembly base for the C ring^14^. The stators are ion-conducting channels and generate torque by converting the electrochemical energy from the ion gradient across the bacterial inner membrane into mechanical energy^14–17^. The flagellar motor can rotate in counterclockwise (CCW) and clockwise (CW) directions, and can switch rapidly between the two rotational states. When all the motors in a bacterial cell rotate in the counterclockwise direction, the filaments form a bundle to propel the cell forward^3,6,7^. Upon activation of the bacterial chemotaxis pathways, the signal protein CheY is phosphorylated by the kinase CheA and binds to the C ring to switch the rotational direction of the motor from counterclockwise to clockwise^2,6,7,18^. Once one or more flagellar motors rotate in the CW state, the filament bundle of the bacterial cell is disassembled, leading to the bacterial cell tumbling and the change of the motility direction^3,6,7^.

The C ring is made up of the proteins FliG, FliM and FliN and attaches to the cytoplasmic face of the MS ring^2,8,9^. The component FliG contains several subdomains. The C-terminal domain of FliG (FliG_C_), consisting of the FliG_CN_ and FliG_CC_ subdomains, interacts with the stator and mediates torque generation and transmission^8,9,19^. The middle domain of FliG (FliG_M_) binds to FliM that also interacts with FliN^8,9^. The N-terminal domain of FliG (FliG_N_) interacts with the C-terminal region of FliF (FliF_C_) and connects the C ring to the MS ring^20^. However, the C ring is highly fragile and easily dissociates from the flagellar motor during biochemical isolation. Despite the fact that a large number of biochemical studies and structural biology efforts have been carried out, the detailed structure, assembly and rotational switching mechanisms of the C ring are still obscure^19–28^. Based on the low-resolution cryo-electron tomography (cryo-ET) analyses of the special flagellar motor in *Borrelia burgdorferi* that harbors the periplasmic flagella, not the general peritrichous or polar flagella, it was proposed that upon the binding of the phosphorylated CheY (CheY-P) to FliM and FliN, FliG tilts outward to interact with the outer rim of the stator, resulting in the protomer elongation and outward expansion of the C ring, which induces the rotational direction switching from CCW to CW^6,25^. However, the outward expansion of the C ring was not observed in other gram-negative bacteria, despite a number of imaging studies^26,27^. Moreover, this model is also not consistent with the findings of the in-frame deletion strain of three residues, Pro-Ala-Ala at positions 169 to 171 of the *Salmonella* FliG, which shortens the sequence, but locks the motor in the CW state^26^. To uncover the rotational switching mechanism, we purified the particles of the intact C ring-containing flagellar motor-hook complex from *Salmonella* Typhimurium and determined two cryo-electron microscopy (cryo-EM) structures of the complex, which are in the default CCW and an active CheY mutant-bound CW states, respectively. The structures reveal that the active mutant CheY protein-induced significant upward movement of the C ring and inward movement of its protomers, not the outward expansion, to mediate the rotational switching, and indicate that the stator units have an unexpected relocation process.

## Results

### Overall structures of the C ring-containing motors in the CCW and CW states

Given the high fragility of the C ring, we examined several reagents as the media of density gradient centrifugation in the purification of the C ring-containing motor-hook complex in the default CCW state from our previously constructed *Λ1 fliCD* strain of *S*. Typhimurium^14^, which lacks the flagellar filament, thus avoiding effects of the filaments on purification of the motor. We successfully obtained particles of the CCW-C ring-containing motor-hook complex for subsequent cryo-EM analysis (Supplementary information, Fig. S1). Most purified particles of the motor-hook complex contain the highly ordered C ring (Fig. 1a, b; Supplementary information, Fig. S1). The cryo-EM density map of the C ring-containing motor-hook complex in the default CCW state was successfully determined to an overall resolution of 7.4 Å (Fig. 1a; Supplementary information, Fig. S1a, S2a, Table S1). Local refinements of the LP ring, MS ring, rod, hook and export apparatus generated their high-quality cryo-EM density maps at the resolutions ranging from 3.0 to 4.3 Å (Supplementary information, Fig. S1a, S2a, Table S1). Initial two-dimensional (2D) classification analyses of manually picked particles revealed that like the MS ring, the C ring adopts a 34-fold symmetric architecture (Supplementary information, Fig. S1a, c). Local refinement of the C ring with C34 symmetry generated a clear density map for the C ring in the CCW state (CCW-C ring) at an overall resolution of 5.9 Å (Fig. 1a; Supplementary information, Fig. S1a, S2a, Table S1). Further local refinement on the four protomers using C34 symmetry-expanded particles improved the density map to a resolution of 4.0 Å (Supplementary information, Fig. S1a, S2). The reconstitution revealed that the structure of the C ring consists of four subrings: the inner, upper, middle and bottom subrings (Fig. 1a, c). The upper, middle and bottom subrings stack in tandem, while the inner subring is relatively separated (Fig. 1a, c). The final structural model of the C ring-containing motor-hook complex in the CCW state contains 341 subunits from 15 proteins, including 34 FliG, 34 FliM, and 102 FliN subunits in the C ring; 34 FliF subunits in the MS ring; 5 FliP, 1 FliR, and 4 FliQ subunits in the export apparatus; 6 FliE, 5 FlgB, 6 FlgC, 5 FlgF, and 24 FlgG subunits in the rod; 26 FlgH and 26 FlgI subunits in the LP ring; and 29 FlgE subunits in the part of the hook (Fig. 1c). The whole structure of the motor-hook complex harbors a molecular weight of ∼10.2 MDa with a height of ∼680 Å, which is much larger than the C ring-free motor (Fig. 1c). The C ring consists of 34 FliG, 34 FliM, and 102 FliN subunits, and has the maximum outer diameter of ∼480 Å at the bottom site (Fig. 1c).

**Fig 1:**
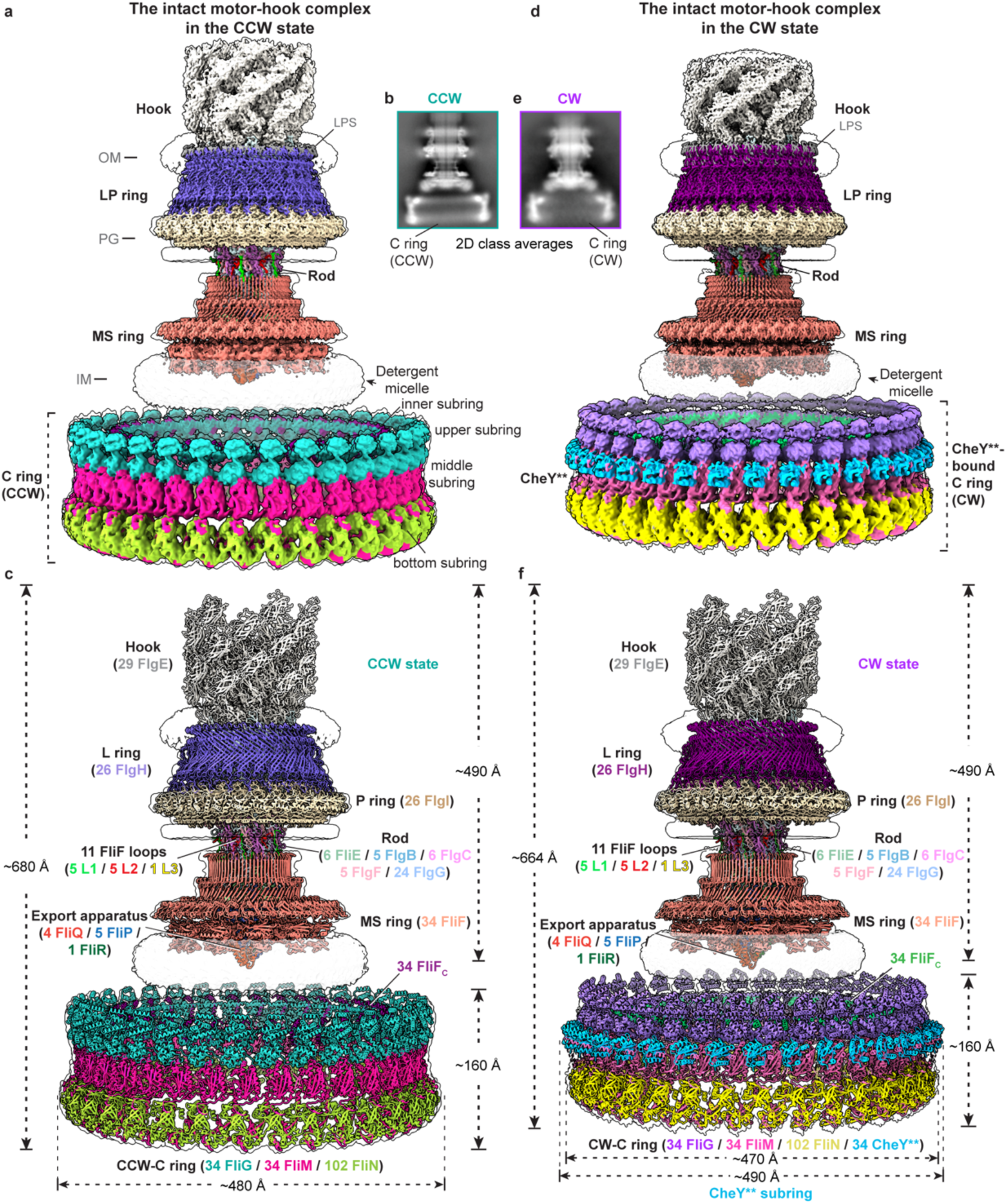
Overall structures of the C ring-containing motor-hook complex in the CCW and CW states. **a**, Composite cryo-EM map of the C ring-containing motor-hook complex in the default CCW state. The map is constructed by integrating six locally refined density maps (the C ring, the distal rod-hook, the LP ring, the proximal rod-export apparatus, the β-collar-RBM3 subrings and the RBM2-RBM1 subrings of the MS ring) into the globally refined 7.4-Å density map of the motor-hook complex in the CCW state. OM, outer membrane; PG, peptidoglycan; IM, inner membrane; LPS, lipopolysaccharide. **b**, Representative 2D class average image of the C ring-containing motor-hook complex in the CCW state. **c**, Overall structure of the C ring-containing motor-hook complex in the CCW state. The subunits in the complex are colored and labeled as indicated. **d**, Composite cryo-EM map of the C ring-containing motor-hook complex in the CheY**-induced CW state. The map is constructed by integrating six locally refined density maps (the C ring, the distal rod-hook, the LP ring, the proximal rod-export apparatus, and the β-collar-RBM3 subrings and RBM2-RBM1 subrings from the MS ring) into the globally refined 4.3-Å density map of the motor-hook complex in the CheY**-induced CW state. **e**, Representative 2D class average image of the C ring-containing motor-hook complex in the CW state. **f**, Overall structure of the C ring-containing motor-hook complex in the CheY**-induced CW state.

To obtain the C ring-containing motor-hook complex in the CW state, we constructed the constitutively active double mutant D13K&Y106W of CheY (hereafter referred to as CheY**), which can mimic CheY-P to tightly bind to the C ring and lock the rotational direction of the motor in the CW state^30,31^. The C ring-containing motor-hook complex in the CW state was obtained by incubating the recombinant CheY** protein with the above purified motor-hook complex in the CCW state. We successfully determined the cryo-EM map of the motor-hook complex in the CheY**-locked CW state to an overall resolution of 4.3 Å (Fig. 1d, e; Supplementary information, Fig. S3, S4a, Table S2). The density maps of the LP ring, MS ring, rod, hook, and export apparatus in the CW state were locally reconstructed to the higher resolutions ranging from 2.8 to 3.8 Å (Supplementary information, Fig. S3a, S4a, Table S2). 2D classification results showed that the C ring in the CW state (CW-C ring) also possess a C34 symmetry (Supplementary information, Fig. S3a, c). The density map of the C ring in the CW state was finally refined to an overall resolution of 5.6 Å with C34 symmetry (Fig. 1d; Supplementary information, Fig. S3a, S4a, Table S2). Further local refinement on four protomers with the symmetry-expanded particles improved the density map to a resolution of 4.4 Å (Supplementary information, Fig. S3a, S4). The subunits of CheY** and the CW-C ring have clear densities in the density map (Fig. 1d; Supplementary information, Fig. S4b). There are in all 34 CheY** subunits, which are bound on the outer surface of the middle subring of the C ring and form a CheY** subring structure (Fig. 1d, f). The final structural model of the CheY**-bound motor-hook complex in the CW state consists of 375 subunits and has a height of ∼664 Å with a maximum diameter of ∼490 Å at the CheY** subring (Fig. 1f). Although the structures of the LP ring, MS ring, rod, hook, and export apparatus in the CW state are highly similar to those of these components in the CCW state (Fig. 1c, f), the protomers of the C ring in the CW state notably undergo inclination and compaction (Fig. 1c, f), suggesting that the binding of CheY** induces significant conformational changes of the C ring for rotational switching.

### Structure of the C ring in the CCW state

The C ring in the CCW state has a height of ∼160 Å with the minimum inner diameters of ∼332 Å at the inner subring, and consists of 34 protomers (Fig. 2a, b; Supplementary information, Fig, S5a). Each protomer is assembled by FliG, FliM and FliN with a stoichiometry of 1:1:3 (Fig. 2c, d). FliG in the CCW-C ring consists of the domains FliG_N_, FliG_M_, FliG_CN_ and FliG_CC_ and adopts a twisted “V”-shaped architecture (Fig. 2e), which is distinct from the extended apo structure of FliG from *A. aeolicus*^19^ (Supplementary information, Fig. S5b). FliM consists of three domains, FliM_N_, FliM_M_ and FliM_C_, and adopts a lifting tong-like structure with FliM_N_ and FliM_C_ as arms (Fig. 2f). FliN is composed of three α-helices and five β-strands, and has a structure similar to FliM_C_ as described^23^ (Fig. 2g). In the C ring, the upper subring is composed of FliG_CC_, while the middle subring is assembled by FliG_CN_, FliG_M_ and FliM_M_ (Fig. 2a-d, h). The bottom subring is formed by FliM_N_, FliM_C_ and FliN (Fig. 2a-d, h). The relatively separate inner subring is constituted by FliG_N_ and FliF_C_ (Fig. 2i).

**Fig 2:**
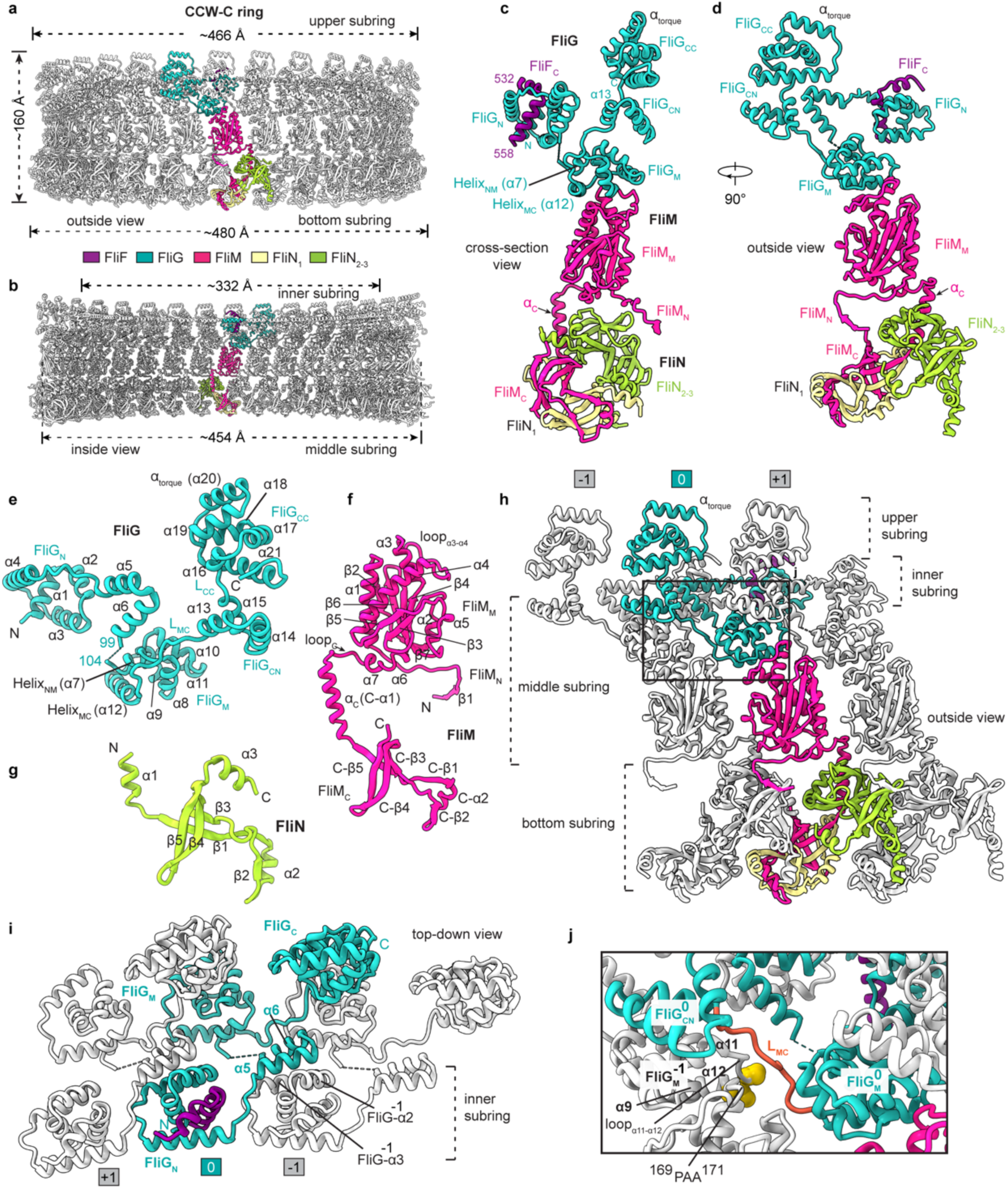
The structure of the C ring in the CCW state. **a-b**, Side (**a**) and cross-section (**b**) views of the structure of the C ring in the CCW state (CCW-C ring). The subunits of FliG, FliM, FliN_1_, FliN_2-3_ and FliF in a protomer are colored in cyan, red, wheat, green and purple, respectively. **c-d**, Cross-section (**c**) and side (**d**) views of the protomer of the CCW-C ring. **e-g**, Structures of the FliG (**e**), FliM (**f**) and FliN (**g**) subunits in the CCW-C ring. **h**, Side view of the inter-protomer interactions in the CCW-C ring. Neighboring protomers are colored in grey and numbered sequentially. **i**. The inter-subunit interactions of FliG in the CCW-C ring. The FliG_N_ domain extends its α5 and α6 helices to interact with the α2 and α3 helices of FliG^-1^ of the prior protomer. **j**, The interactions of the L_MC_ loop with FliG_M_^-1^ in the CCW-C ring. The residues P169, A170 and A171 of FliG_M_^-1^ are shown as spheres in yellow.

The protomer of the C ring has a “Y”-shaped architecture (Fig. 2c, d), and obliquely binds to the neighboring two protomers in a “S”-like manner (Fig. 2h), which generates extensive inter-protomer interactions within the C ring. In one protomer, the helices α5 and α6 of FliG_N_ pack onto helices α2 and α3 of the FliG^-1^ subunit of the prior protomer. A flexible loop between α6 and Helix_NM_ (α7), which is disordered in the structure, links FliG_N_ to the helix α7 of FliG_M_ (Fig. 2e, i). FliG_M_ binds to the upper surface of FliM_M_ via the α9, α11 and Helix_MC_ (α12) helices and the loop_α8-α9_ (Fig. 2c, 2e; Supplementary information, Fig. S5c). The following FliG_CN_ domain extends outwards and binds onto the upper surface of the FliG_M_ domain in the prior protomer (Fig. 2h, j). The long linking loop L_MC_, which connects FliG_M_ and FliG_CN,_ binds into a cleft between α9 and loop_α11-α12_ of the FliG_M_^-1^ domain, close to the residues P169-A170-A171 (^169^PAA^171^) (Fig. 2j), the deletion of which caused the CW-biased rotation of the motor^32^, suggesting that the interactions play a key role in the rotational switching. The α15 helix of FliG_CN_ and the following hinge loop L_CC_, which is formed by the conserved M_233_FLF_236_ motif^9^, brace the FliG_CC_ domain via hydrophobic interactions and stabilize its orientation in the upper region of the protomer (Supplementary information, Fig. S5d). The α_torque_ (α20) helix of FliG_CC_, which interacts with the MotA subunits of a stator unit in torque transmission^9^, lies on the top of the C ring (Fig. 2h; Supplementary information, Fig. S5e). The stator-interacting charged residues, R281 and D288/D289^33^ are located at the N- and C-terminus of α_torque_, respectively (Supplementary information, Fig. S5e). The head-to-tail packing of the α_torque_ helices in the upper subring generates a periodic distribution of positive-to-negative charges along the CCW direction on the upper surface of the C ring (Supplementary information, Fig. S5f).

### The inter-subunit interactions of the C ring in the CCW state

The FliM and FliN subunits are extensively involved in the intra- and inter-protomer interactions in the C ring. FliM_M_ interacts with the prior and next FliM_M_ domains in the middle subring in a “shoulder-to-shoulder” manner via α5, loop_α3-α4_, α1, α_C_, and loop_C_ (Fig. 3a, b). The interfaces generate the relatively weak interactions and leave small gaps between the FliM_M_ domains (Fig. 3a). FliM_M_ also contacts α8 of the FliG_M_^+1^ domain of the next protomer via loop_α1-β2_ (Fig. 3a, c) and interacts with α_C_ and loop_C_ of the prior FliM_C_ domain to strengthen the middle subring and bridge the middle and bottom subrings, respectively (Fig. 3a, d).

**Fig 3:**
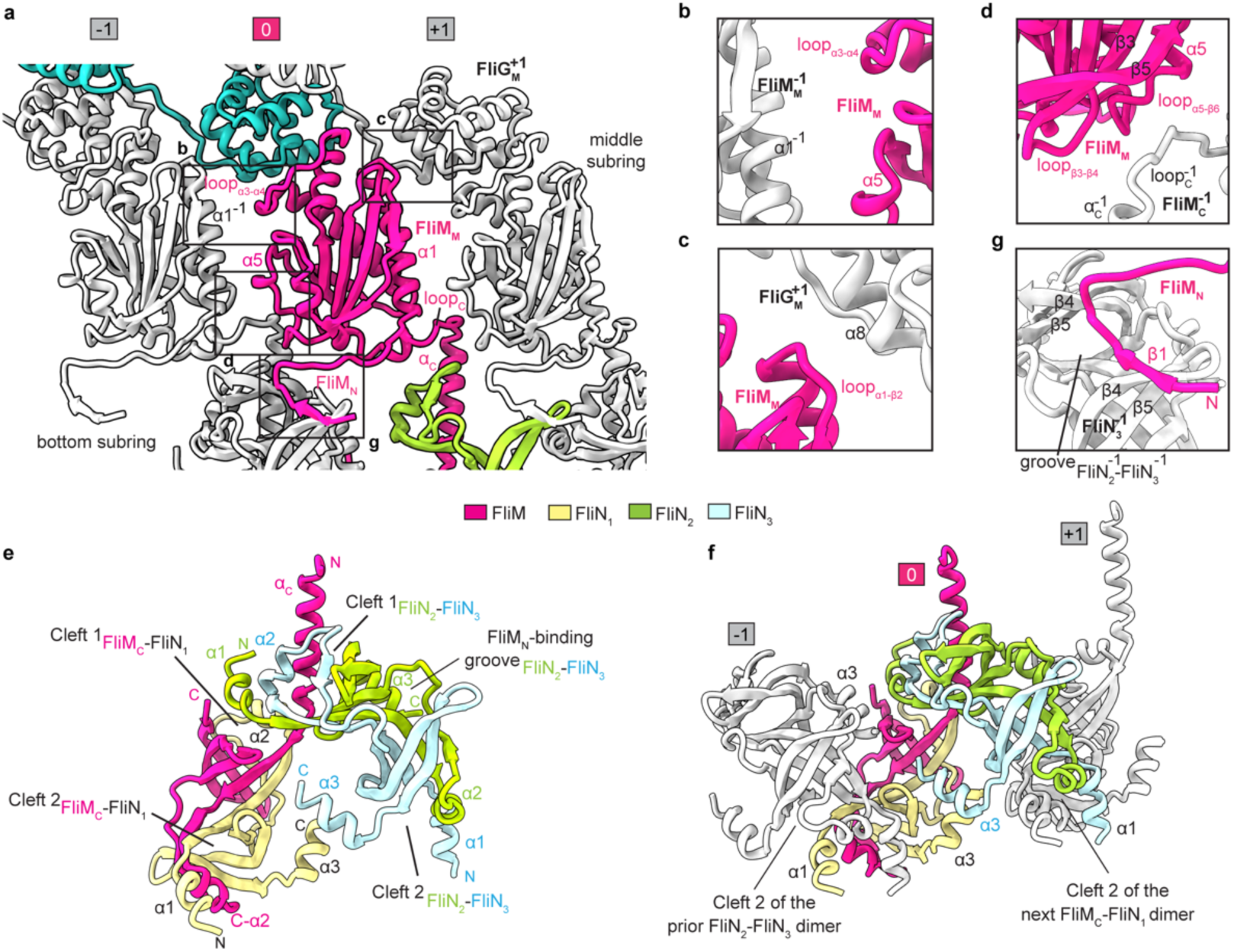
The inter-subunit interactions of the C ring in the CCW state. **a-d**, The inter-protomer interactions mediated by FliM_M_ in the CCW-C ring. Neighboring FliM_M_, FliM_N_, and FliG_M_ domains are colored in grey and numbered sequentially (**a**). Detailed interactions of the FliM_M_ with neighboring FliM_M_^-1^ (**b**), FliG_M_^+1^ (**c**) and FliM_C_^-1^ (**d**) domains are illustrated as indicated. **e**, Side view of the FliM_C_-FliN tetramer in the CCW-C ring. **f**, The spiral structure of the bottom subring of the CCW-C ring and the α1-swapping interactions through Cleft 2 of the FliM-FliN tetramers between the protomers. **g**, Interactions of FliM_N_ with the FliN_2_-FliN_3_ dimer of the prior protomer. The FliM, FliN_1_, FliN_2_ and FliN_3_ subunits of the focused protomer are colored in red, wheat, green and light blue respectively in (**a**), (**e**) and (**f**). The FliM and its interacting domains are colored in red and grey in (**b-d**) and (**g**).

In the bottom subring, FliM_C_ forms a heterodimer with the FliN_1_ subunit in a head-to-tail manner (Supplementary information, Fig. S5g), while the FliN_2_ and FliN_3_ subunits form a homodimer in the same fashion (Supplementary information, Fig. S5h). The dimerization generates two symmetrical clefts (Cleft 1 and Cleft 2) at each end of the dimers (Supplementary information, Fig. S5g, h). The two dimers undergo further dimerization and form a tetrameric structure through swapping the binding of α_C_ of FliM_C_ and α1 of FliN_2_ into Cleft 1 of each other in the protomer (Fig. 3e; Supplementary information, Fig. S5i). The tetramerization is also strengthened by the interactions between the α3 helices of FliN_1_ and FliN_3_ (Fig. 3e; Supplementary information, Fig. S5i). The α1 helix of FliN_1_ extends to bind into Cleft 2 of the FliN_2_-FliN_3_ dimer in the prior protomer, while the α1 of FliN_3_ extends to bind into Cleft 2 of the FliM_C_-FliN_1_ heterodimer in the next protomer (Fig. 3f). The α1-swapping interactions across Cleft 2 between the protomers generate a continuous spiral structure for the bottom subring with the FliM_C_-FliN_1_ dimer as the inner side and the FliN_2_-FliN_3_ dimer as the outer side (Fig. 3f; Supplementary information, Fig. S5j, k). The N-terminal region (residues 34-45) of FliM_N_ binds to the central groove in the prior FliN_2_-FliN_3_ dimer via the formation of an antiparallel β-sheet with β4 of FliN_3_^-1^ to lift up the outer side of the bottom subring (Fig. 3a, g).

### The structure of the C ring in the CW state

The CheY**-bound C ring in the CW state has a height of ∼160 Å, which is the same as that of the C ring in the CCW state (Fig. 4a). However, the upper, middle and bottom subrings of the CW-C ring have diameters of ∼446 Å, ∼440 Å and ∼470 Å, respectively (Fig. 4a, b), which are all smaller than those in the CCW-C ring, indicating that the binding of CheY** induces a more compressed structure of the C ring. The inner diameter of the inner subring is also shortened to ∼312 Å (Fig. 4b). In the CW-C ring, FliG_C_ adopts a conformation that differs from that of the crystal structure of FliG_C_ in the ^169^PAA^171^ mutant of FliG^22^ (Supplementary information, Fig. S6a). Structural superimposition revealed that related to FliG_M_, the FliG_N_ domain of FliG in the CW-C ring rotates by ∼140° to interact with the helices α2 and α3 of the FliG_N_ domain of the next protomer (Supplementary information, Fig. S6b, c). CheY** is bound into an open groove between two adjacent protomers on the surface of the middle subring of the C ring and interacts with FliG_M_, FliM_M_ and the prior FliM_M_ domain (Fig. 4c-e). But in contrast to previous predictions^34^, CheY**has no interactions with FliN and the bottom subring in the structure.

**Fig 4:**
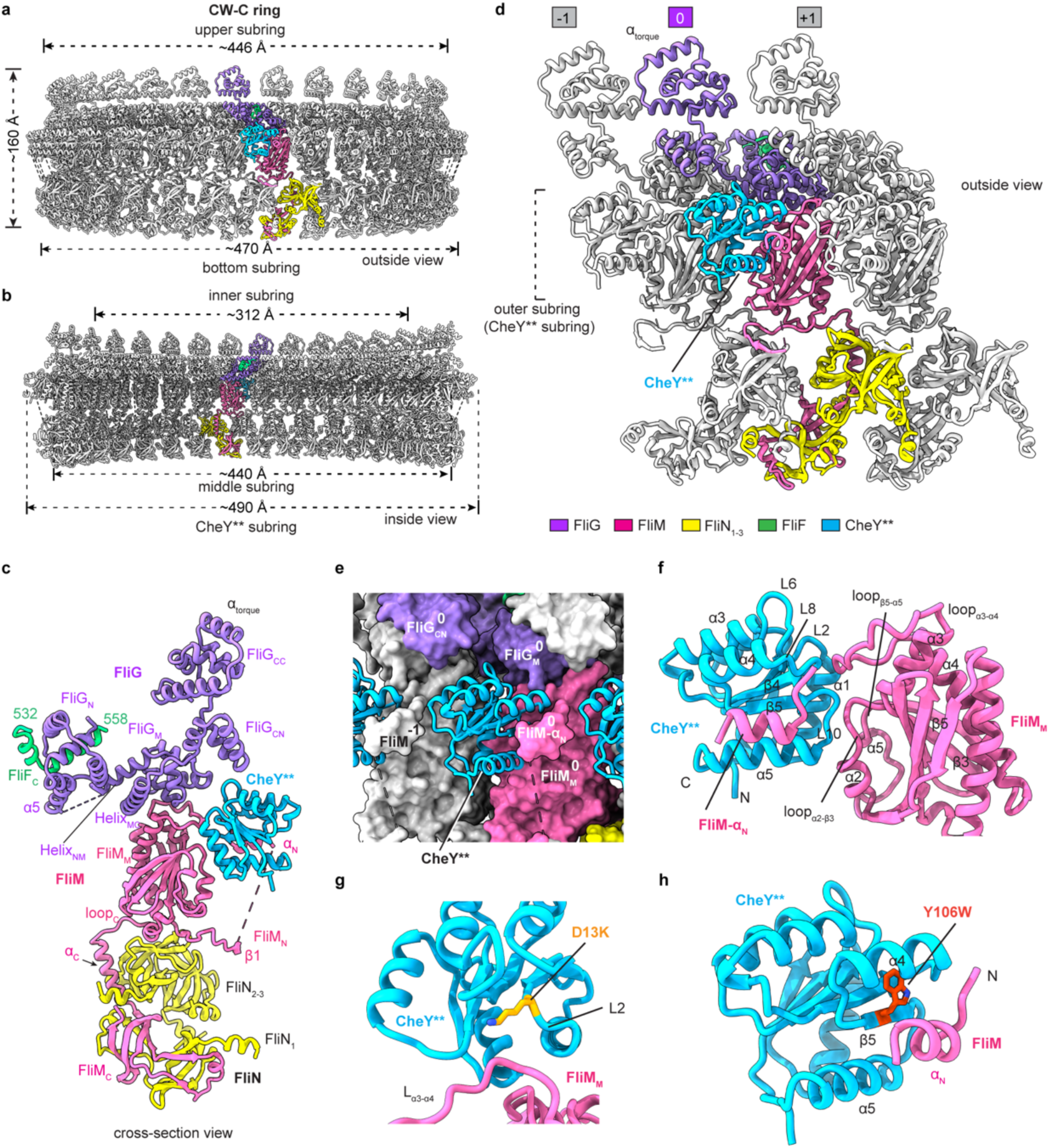
The structure of the C ring in the CW state. **a-b**, Side (**a**) and cross-section (**b**) views of the structure of the C ring in the CheY**-induced CW state (CW-C ring). The subunits of FliG, FliM, FliN, FliF_C_ and CheY** in a protomer are colored in medium purple, pink, yellow, green and blue, respectively. **c**, Cross-section view of the protomer of the CW-C ring. **d**, Inter-protomer interactions of the CW-C ring. Neighboring protomers are colored in grey and numbered sequentially. **e**, Surface representation of the interactions of CheY** with the FliG and FliM domains of the CW-C ring. CheY** is shown as cartoon. **f**, Detailed interactions of CheY** with FliM_M_. **g-h**, The interactions of the mutant residues D13K (**g**) and Y106W (**h**) of CheY** with FliM.

The structure of CheY** is highly similar to that of the beryllium fluoride (BeF3^-^)-activated CheY^35^ (Supplementary information, Fig. S6d). At the interface with FliM, CheY** interacts with FliM_M_ and the N-terminal loop and the α_N_ helix of FliM_N_ (Fig. 4f). The α1 helix, L2, L6, L8 and L10 loops of CheY** form an open concave surface to interact with loop_α2-β3_, loop_α3-α4_, loop_β5-α5_ and α5 of FliM_M_ (Fig. 4f). CheY** binds to α_N_ and the N-terminal loop of FliM_N_ via the α4-β5-α5 surface (Fig. 4f). The mutated residue D13K of CheY** interacts with the loop_α3-α4_ of FliM_M_ (Fig. 4g), while the mutated residue Y106W is located on the α4-β5-α5 surface of CheY** and stacks with the α_N_ helix of FliM_N_ (Fig. 4h), suggesting that the double mutation activates CheY through generating direct interactions with the C ring. In the interactions with FliG, the L6 and L8 loops of CheY** clamp the α11 helix of FliG_M_ (Supplementary information, Fig. S6e). CheY** also contacts with the β2, β6 and β7 strands and the linking loop_β6-β7_ of FliM_M_^-1^ of the prior protomer via its α1-β2 surface (Supplementary information, Fig. S6f), which enhances the inter-protomer interactions in the middle subring.

### The CheY-induced conformational changes of the C ring

Structural and cryo-EM map superimpositions of the C ring-containing motor-hook complexes in the CW and CCW states reveal that upon the binding of CheY**, the whole structure of the C ring in the CW state undergoes a significant upward shift of ∼16 Å towards the MS ring and the bacterial inner membrane (Fig. 5a; Supplementary information, Fig. S7a), which reduces the distance between the MS ring and C ring and shortens the overall length of the motor-hook complex (Fig. 1c, f). Structural comparison of the CCW- and CW-C rings revealed that, in contrast to the previous assumption of the outward expansion of the C ring for the rotational switching^6,25^, all four subrings of the CW-C ring undergo significant inward contractions towards the center of the motor upon the binding of CheY**, resulting in the smaller diameters of the four subrings (Fig. 5b; Supplementary information, Fig. S7b). The inward contraction of the inner FliG_N_-FliF_C_ subring decreases its inner diameter by ∼20 Å, while the outer diameter of the bottom subring is shortened by ∼10 Å (Fig. 5b).

**Fig 5:**
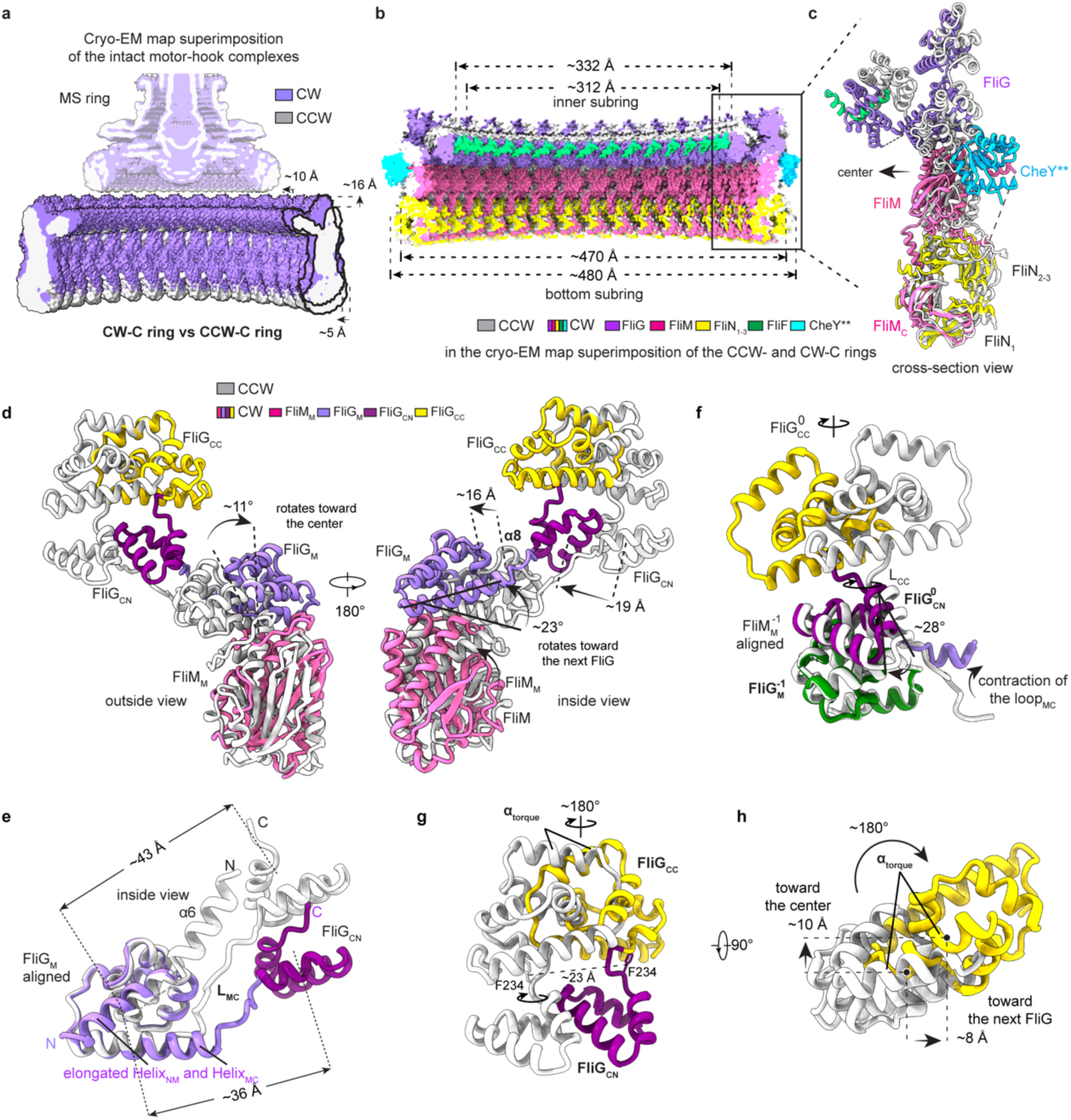
The CheY**-induced conformational changes of the C ring in the intact motor. **a**, Cryo-EM map superimposition of the C ring-containing motor-hook complexes in the CCW and CW states. The density maps of the motor-hook complexes in the CCW and CW states are colored in grey and medium purple, respectively. The changes are indicated by black arrows. **b**, Structural comparison of the CCW- and CW-C rings in the cryo-EM map superimposition of the CCW- and CW-C rings in Fig **S7b**. **c**, Structural comparison of the protomers of the CCW- and CW-C rings in the cryo-EM map superimposition of the C rings in (**b**). The protomers in the CCW- and CW-C rings are colored in grey and as in Fig 4a, respectively. **d**, Structural comparison of the FliG_M_-FliG_C_-FliM_M_ domains in the structural superimposition of the CCW- and CW-C-ring structures. FliM_M_, FliG_M_, FliG_CN_ and FliG_CC_ from the CW-C ring are colored in pink, medium purple, purple and yellow, respectively. The domains in the CCW-C ring are colored in grey. Conformational changes are indicated by black arrows. **e**, Structural comparison of the Helix_NM_ and Helix_MC_ helices from the CCW- and CW-C rings through structural superimposition of FliG via the FliG_M_ domains. The elongated Helix_NM_ and Helix_MC_ in the FliG subunit are labeled as indicated. **f**, Structural comparison of the FliG_CN_-FliG_CC_ domains in the CCW- and CW-C rings by superimposing the FliG_M_^-1^ domains. **g-h**, Side (**g**) and top (**h**) views of the conformational changes of the FliG_CC_ domain in the structural superimposition of the CCW- and CW-C rings. The residues of F234 are shown as sticks and labeled as indicated (**g**).

In addition to the overall upward movement of the C ring in the motor, the binding of CheY** remodels the structures of the protomers in the CW-C ring (Fig. 5c). With the central β-strands of FliM_C_ as the fulcrum, FliM_M_ and α_C_ of FliM_C_ obliquely rotate by ∼11° towards the center of the C ring, leading to the inward movement of the N-terminus of α_C_ of FliM_C_ by ∼11 Å and of the FliG_M_-binding loop_α3-α4_ of FliM_M_ by ∼21 Å (Supplementary information, Fig. S7c). The FliN_2_-FliN_3_ dimer also undergoes a rotation by ∼11°, thereby retaining the inter-subunit interactions between FliM_C_ and the FliN subunits in the bottom subring (Supplementary information, Fig. S7d). In addition, FliM_M_ tilts by ∼23° to the FliM_M_ domain of the next protomer (Supplementary information, Fig. S7e). The loop_C_ moves ∼8 Å towards the center of the C ring and has a more extended conformation, thereby reducing the contacts of loop_C_ to the next FliM_M_ domain (Supplementary information, Fig. S7e). The N-terminal region of FliM_N_, which binds the FliN_2_-FliN_3_ dimer in the prior protomer, moves ∼11 Å towards the center of the C ring (Supplementary information, Fig. S7f).

The binding of CheY** onto the C ring not only changes the orientations of all the domains of FliG, but also alters the distances between the domains in the protomer, leading to the overall structure of FliG in the CW state to be distinct from that in the CCW state (Supplementary information, Fig. S7g). The packing of FliG_N_, FliG_M_ and FliG_C_ in FliG in the CW state is more compacted than that in the CCW state (Supplementary information, Fig. S7g). In company with the rotation of FliM_M_, FliG_M_ undergoes a rotation by ∼11° towards the center of C ring and a tilt by ∼ 23° toward the next protomer, leading to a movement by ∼16 Å of the upper region of the α8 helix that is involved in the interactions with the FliG_CN_^+1^ domain of the next protomer (Fig. 5d). The FliG_CN_ domain also moves by ∼19 Å towards the center of the C ring (Fig. 5d). Despite the conformational changes, FliG_M_ in the CW state maintains an identical interface with FliM_M_ as that in the CCW state (Fig. 5d). FliG_N_ moves downwards by ∼25 Å and rotates by ∼180° towards the prior protomer (Supplementary information, Fig. S7h), resulting in the reduced diameter of the inner subring (Fig. 5b). The conformational changes of FliG_N_ induces the fusion of the α6 helix of FliG_N_ with the N-termini of the Helix_NM_ helix of FliG_M_, which generates a longer helical structure for Helix_NM_ of FliG_M_ in the CW-C ring (Fig. 5e).

### The conformational changes of FliG_CC_ for rotational switching

The binding of CheY** also induces a significant contraction of the loop L_MC_ that connects FliG_M_ and FliG_CN_ and results in the formation of an elongated Helix_MC_, which not only reduces the distance between FliG_M_ and FliG_CN_ by ∼7 Å (Fig. 5e), but also causes the clockwise rotation of FliG_CN_ by ∼28° on the binding surface of the FliG_M_^-1^ domain in the prior protomer (Fig. 5f; Supplementary information, Fig. S8a). The rotation and movement of FliG_CN_ cause the rotation by ∼180° and relocation of the hinge loop L_CC_ that connects FliG_CN_ and FliG_CC_ (Supplementary information, Fig. S8b). F234, the key residue stabilizing the orientation of FliG_CC_ in the protomer via hydrophobic interactions, in L_CC_ moves by ∼23 Å and rotates by 180° (Fig. 5g; Supplementary information, Fig. S8b). As the consequence, the whole FliG_CC_ domain undergoes the rotation by ∼180° and movements by ∼8 Å toward the next FliG_CC_ domain and inward contraction by ∼10 Å to the center of the C ring (Fig. 5g, h; Supplementary information, Fig. S8c, d), which results in a reversed orientation of the α_torque_ helix and changes the periodic CCW distribution of the charged residues R281 and D288/D289 to be a CW distribution on the upper surface of the CW-C ring (Fig. 6a). The changed distribution of the charge residues then reverses the electrostatic surface potential of the α_torque_ in the CW-C ring (Supplementary information, Fig. S5f, S8e). Thus, in the intact flagellar motor, the binding of CheY** onto the C ring finally induces the upward shift by ∼16 Å, an inward movement by ∼10 Å and the orientation reversion of α_torque_ to trigger the rotational direction switching of the motor from CCW to CW (Fig. 5g, h, 6b).

**Fig 6:**
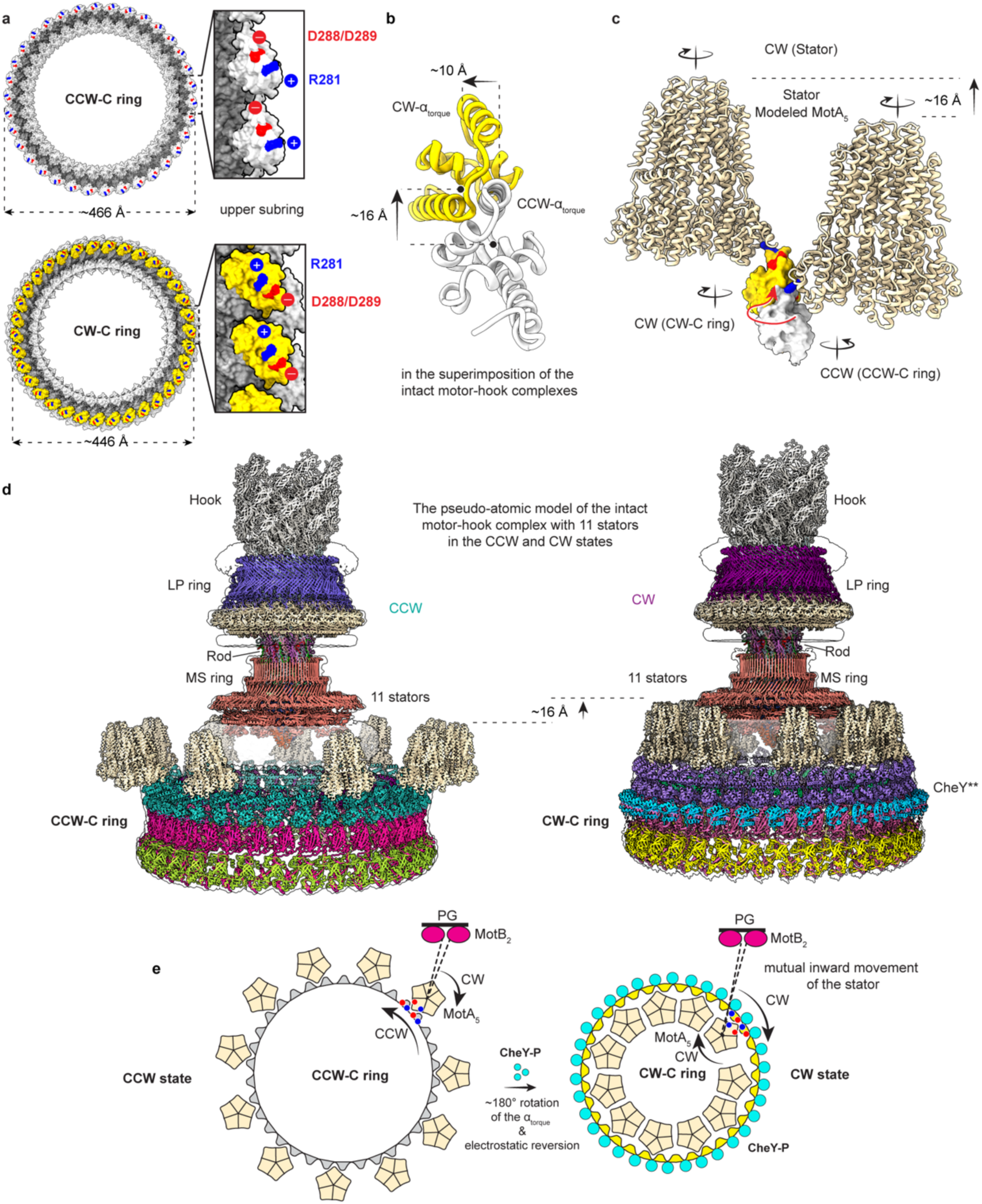
The proposed rotational switching mechanism of the bacterial flagellar motor. **a**, Distribution of the charged residues R281, D288 and D289 on the upper surface of the CCW-C ring (upper) and the CW-C ring (bottom). The charged residues of the α_torque_ helices are colored in blue and red as positive and negative charges, respectively. The FliG_CC_ domains of the CCW- and CW-C rings are colored in grey and yellow, respectively. **b**, Cross-section view of the conformational changes of the FliG_CC_ domain in the cryo-EM map superposition of the motor-hook complexes. In addition to the conformational changes illustrated in Fig 5h, the FliG_CC_ domain shifts upward by ∼16 Å in the motor in the CW state. **c**, Side view of the relocation of a stator unit in the motor for the rotational switching. The structural models of the interactions of the MotA_5_ pentamer of a stator with FliG_CC_ were constructed according to the cryo-ET map of the *Helicobacter* motor in the CCW state^36^, as shown as in Fig **S9a**. The stator units are colored in wheat and shown as cartoons. The FliG_CC_ domains are colored as shown in (**a**) and are shown as surface. **d,** The pseudo-atomic models of the intact motor-hook complex with 11 stators in the CCW (left) and CW (right) states. **e**, Schematic diagram of the rotational switching mechanism of the flagellar motor. The binding of CheY** or CheY-P onto the C ring induces the upward and inward movements of the FliG_CC_ domains and reverses the orientations of the α_torque_ helices, which induces the relocation of the stator units in the bacterial inner membrane from the outer side (left) of the upper subring to the inner side (right) and triggers the rotation switching from CCW to CW.

The CheY**-induced upward movement and inward contraction of the C ring suggest that the stator units need to be relocated in the bacterial inner membrane through inward movements by an appropriate distance towards the center of the motor for the rotational switching. A 24-Å cryo-ET map of the *Helicobacter* flagellar motor in the CCW state^36^ clearly shows that the stator units are bound onto the top surface of the C ring at the outer side of the upper subring (Supplementary information, Fig. S9a). The structural models of the protomer of the *Salmonella* CCW-C ring and of the MotA pentamer (MotA_5_) of a stator unit could be well matched to the densities of the *Helicobacter* C ring and the stators, respectively, in the low-resolution cryo-ET map^36^ (Supplementary information, Fig. S9a). For each stator unit in the map, two adjacent FliG_CC_ domains of the C ring clamp the protruding MotA subunit from its two sides (Supplementary information, Fig. S9a). According to the interactions of the protomers with the stator, a pseudo-atomic structural model of the intact *Salmonella* flagellar motor with stator units in the CCW state was modeled (Fig. 6c, d; Supplementary information, Fig. S9b), The interactions of the protomers with a stator unit only allowed 11 stator units as the maximum number to be bound onto the C ring in the *Salmonella* motor, which is consistent with the previous biophysical analyses^37,38^ (Fig. 6d; Supplementary information, Fig. S9b). The structural model of the intact motor with the 11 stator units was perfectly matched to the ∼69 Å cryo-ET maps of the *Salmonella* flagellar motor in the CCW state applied with C11 symmetry^39^ (Supplementary information, Fig. S9c) . The pseudo-atomic structural model of the *Salmonella* motor with the 11 stator units in the CW state was then built, according to the conformational changes of the FliG_CC_ domains (Fig. 6c, d; Supplementary information, Fig. S9b). The relocated stators did not generate any structural collision and did not allow accommodation of more stator units on the C ring either (Fig. 6d; Supplementary information, Fig. S9b). We modeled the structure of the CW-C ring-containing motor-hook complex of *S*. Typhimurium into the ∼19-Å cryo-ET density map of the flagellar motor of *Vibrio alginolyticus* in a FliG-G215A mutant-based biased CW state^27^ (Supplementary information, Fig. S9d). The structure of the *Salmonella* C ring is well fitted with the cryo-ET density map of the *Vibrio* flagellar motor that also contains 34 protomers in the C ring^27^ (Supplementary information, Fig. S9d). The stator units in the CW state could be well accommodated in the notably protruding densities above the C ring in the inner membrane, which are closed to the inner side of the upper subring and were not annotated previously in the map^27^ (Supplementary information, Fig. S9d), suggesting that the stator units on the *Vibrio* motor indeed undergo inward relocation in the inner membrane. Thus, the conformational changes of the CW-C ring indicate that the stator units require a relocation process for adaptation with FliG_CC_ during the rotational switching (Fig. 6d, e; Supplementary information, Fig. S9b).

## Discussion

It is a long-standing question in the field about the rotational switching mechanism of the bacterial flagellar motor^2,3,6,7^. The two cryo-EM structures of the *Salmonella* flagellar motor-hook complexes in the CCW and CW states, respectively, not only present the intact architectures of a flagellar motor, but also uncover the detailed structures and assembly of the CCW-ad CW-C rings. In contrast to the previously reported or proposed structures^19^, FliG adopts a twisted “V”-shaped structure in the CCW-C ring and a more compacted structure in the CW-C ring. CheY** not only binds to FliM, but also interacts with FliG and the FliM^-1^ subunit in the prior protomer in the middle subring of the C ring. The binding of CheY** does not cause any outward expansion of FliG or the C ring. Instead, the C ring undergoes a notable upward movement toward the MS ring and the bacterial inner membrane, and inward contraction to its center. The upward movement of the C ring likely promotes the relocation and accommodation of the stator units in the inner membrane around the motor. The CheY**-induced conformational changes of Helix_MC_ and L_MC_ and the subsequent structural changes of FliG_CN_ and L_CC_ eventually reverse the orientation of α_torque_ and electrostatic distribution on the upper surface of the C ring to trigger the rotational direction switching of the motor (Fig. 6e).

The C ring plays a key role in the mechanosensing of the bacterial flagellar motor to mechanical stimuli in environments by mediating the loading of the stator units^5,13,38,40^. The relatively separated FliG_N_-FliF_C_ subcomplex is connected to FliG_M_ via Helix_NM_ that tightly stacks with Helix_MC_ in the C ring. It is possible that the C ring adjusts the conformations of the FliG_CC_ domains via the Helix_NM_-Helix_MC_-FliG_CN_-FliG_CC_ module cascade upon the conformational changes of the FliG_N_-FliF_C_ subcomplexes and the MS ring under environmental stresses to load the stators onto the motor (Supplementary information, Fig. S10a). In the C ring-containing motor-hook complexes, the MS ring is more stable and harbors much more clear densities for the outer RBM1 subring, outer RBM2 subring and putative inner RBM1 subring^41,42^, which are made up of 11 RBM1, 11 RBM2 and 23 RBM1 domains of FliF, respectively (Supplementary information, Fig. S10b-d), suggesting that the C ring is also important for the stability of the MS ring. Moreover, besides the 5 L1 and 5 L2 loops, there is one more peptide loop L3, which consists of the residues G310-P322 of FliF and is also extended from the β-collar region of the MS ring, bound in a hydrophobic groove that is formed by the α2-α3 linking loop of FliE^6^ with the D0 and D_C_ domains of FlgC^1^ and FlgC^6^ on the surface of the rod (Supplementary information, Fig. S10e-g). The interaction manner of L3 with the rod was not observed in the interactions of L1 and L2 loops with the rod in the structure of the C ring-free flagellar motor^14^. It is possible that the 11 loops are derived from the FliF subunits in the 11-fold outer subrings. The extensive inter-promoter interactions between the “Y”-shaped protomers in the C ring suggest that the conformational changes of the protomers have synergistic effects on recruiting CheY-P for rotational switching^43^ (Supplementary information, Fig. S10a).

It was assumed that the stator units were fixed in their positions during the rotational switching^7,25,28^. The structures of the C ring-containing motor-hook complexes and the structural modeling of the intact motors with the stators in the low-resolution cryo-ET maps reveal that the stator units undergo a relocation adaptation process in accompany with the CheY-induced conformational changes of the C ring. In the CCW state, the stators, which rotate in the CW direction, are located at the outer side of the upper subring of the C ring and trigger the rotation of the motor in the CCW direction. Upon the binding of CheY-P, the stator units are relocated to the inner side of the upper subring of the C ring for adaptation with the reversed orientation of the α_torque_ helix of FliG_CC_, leading to the CW-direction rotation of the motor (Fig. 6e; Supplementary information, Fig. S9b). For the relocation towards the center of the motor, the stator units probably undergo rotation by 180° and inward movements in the bacterial inner membrane. There is another possibility that the bacterial inner membrane is lift up by the MS ring and CW-C ring or undergoes inward movement around the flagellar motor, leading the relocation of the stator units. Despite the diversities of the flagellar motors, the motor components of *S*. Typhimurium are highly conserved in bacterial species^44^. The inward relocation of the stators is possible a general adaptation process for the rotational switching of the flagellar motors.

## Data availability

Cryo-EM density maps and coordinates related to the flagellar motor-hook complex in the CCW state have been deposited in EMDB and PDB under the following accession codes: the flagellar C ring-containing motor-hook complex (EMD-37679, PDB ID: 8WO5); the protomers of the CCW-C ring (EMD-38546, PDB ID: 8XP0); the CCW-C ring with C34 symmetry (EMD-39349, PDB ID: 8YJT); the membrane-anchored part, including the rod, export apparatus, LP ring, MS ring and hook, of the motor-hook complex in the CCW state (EMD-37630, PDB ID: 8WLT); the LP ring with C26 symmetry (EMD-37618, PDB ID: 8WLE); the whole MS ring with the proximal rod and export apparatus (EMD-37625, PDB ID: 8WLN); the whole rod with the export apparatus and the part of the hook (EMD-37628, PDB ID: 8WLQ); the distal rod with the part of the hook (EMD-37627, PDB ID: 8WLP); the proximal rod with the export apparatus and 11 FliF loops (EMD-37619, PDB ID: 8WLH) and the β-collar-RBM3 subrings of the MS ring with C34 symmetry (EMD-37620, PDB ID: 8WLI). Cryo-EM density maps and coordinates related to the flagellar motor-hook complex in the CW state have been deposited in EMDB and PDB under the following accession codes: the intact CheY**-bound motor-hook complex (EMD-37684, PDB ID: 8WOE); the protomers of the CW-C ring (EMD-38547, PDB ID: 8XP1); the CheY**-bound CW-C ring with C34 symmetry (EMD-37570, PDB ID: 8WIW); the membrane-anchored part, including the rod, export apparatus, LP ring, MS ring and hook, of the motor-hook complex in the CW state (EMD-37611, PDB ID: 8WL2), the LP ring with C26 symmetry (EMD-37547, PDB ID: 8WHT); the whole MS ring with the proximal rod and export apparatus (EMD-37605, PDB ID: 8WKQ), the whole rod with the export apparatus and the part of the hook (EMD-37601, PDB ID: 8WKK); the distal rod with the part of the hook (EMD-37600, PDB ID: 8WKI); the proximal rod with the export apparatus and 11 FliF loops (EMD-37594, PDB ID: 8WK3); the β-collar-RBM3 subrings of the MS ring with C34 symmetry (EMD-37590, PDB ID: 8WJR) and the β-collar-RBM3 subrings of the MS ring with the proximal rod (EMD-37595, PDB ID: 8WK4).

## Acknowledgements

We thank the core facility of Life Sciences Institute Zhejiang University for equipment support and the cryo-EM centers of Zhejiang University, Fudan University, Institute of Precision Medicine of Shanghai Jiaotong University School of Medicine for their assistance in cryo-EM data collection. This work was supported by grants from NSFC (U23A20163 and 81925024), the National Key Research and Development Program of China (2017YFA0503900), and the Fundamental Research Funds for the Central Universities. Y. Zhu and Y. Zhou were supported by National High-level Talents Special Support Program of China.

## Author contribution

Y. Zhu conceived this study. Y. Zhu and Y. Zhou supervised the study; J.T., L.Z. and Y. Zhu designed experiments; J.T. and L.Z purified the complexes and determined cryo-EM structures; X.Z. and S. H. assisted assays; J.T. and Y. Zhu prepared the manuscript.

## Competing interests

The authors declare no competing interests.

## Corresponding author

Correspondence to Y. Zhu, Y. Zhou

## Supplementary information

**Supplementary information, Figure S1.**
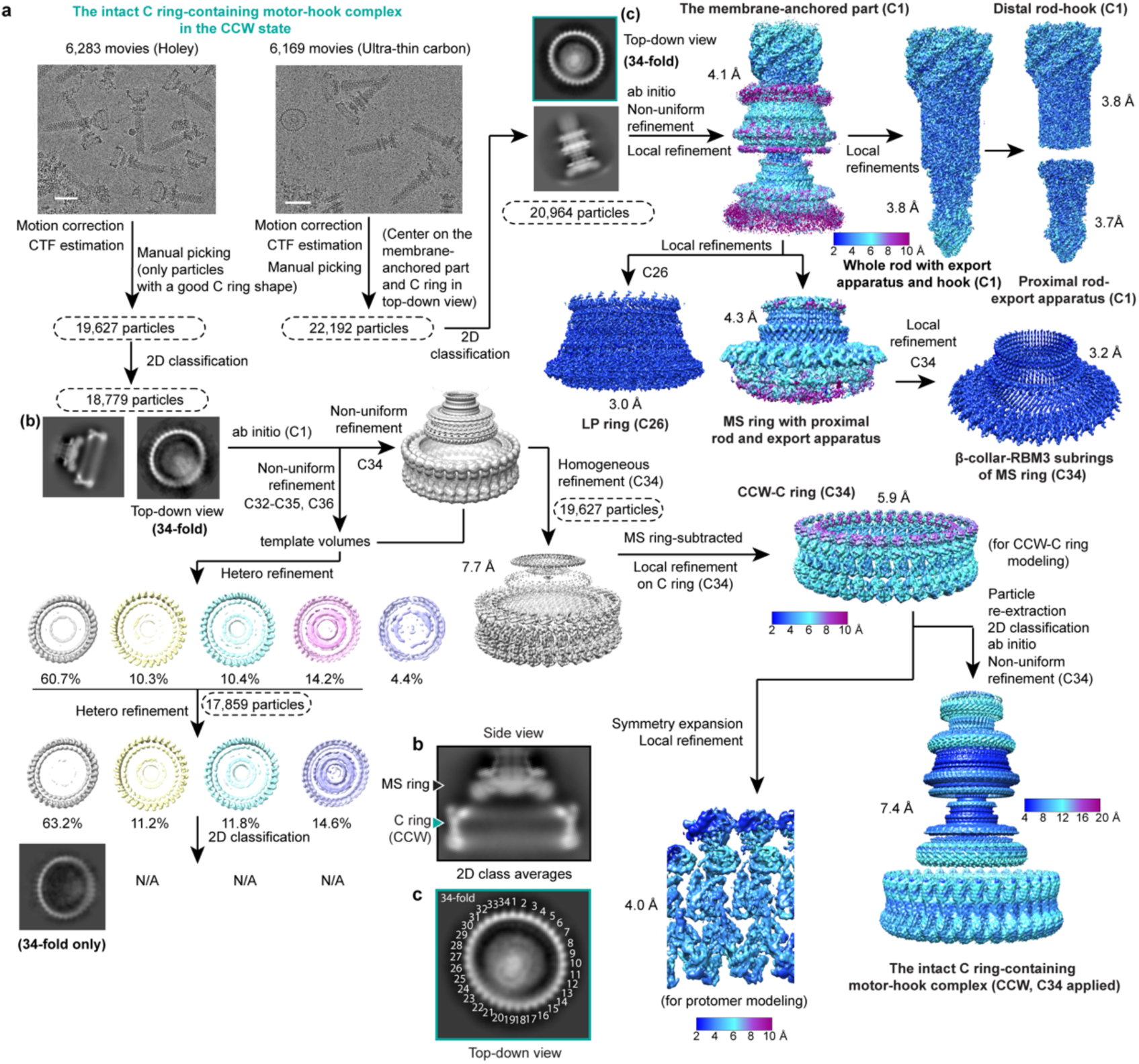
Cryo-EM data processing of the flagellar motor-hook complex in the CCW state. **a**, The flow chart for the cryo-EM data processing of the C ring-containing motor-hook complex in the CCW state. The local resolution maps were calculated in cryoSPARC using two independent half maps of each reconstruction as input. All density maps were prepared using *Chimera* and *ChimeraX*. Scale bar for the cryo-EM micrographs, 50 nm. N/A, not available. **b**, A representative 2D class average of the C ring in the CCW state with the MS ring from the side view. **c**, A representative 2D class average of the CCW-C ring from the top-down view.

**Supplementary information, Figure S2.**
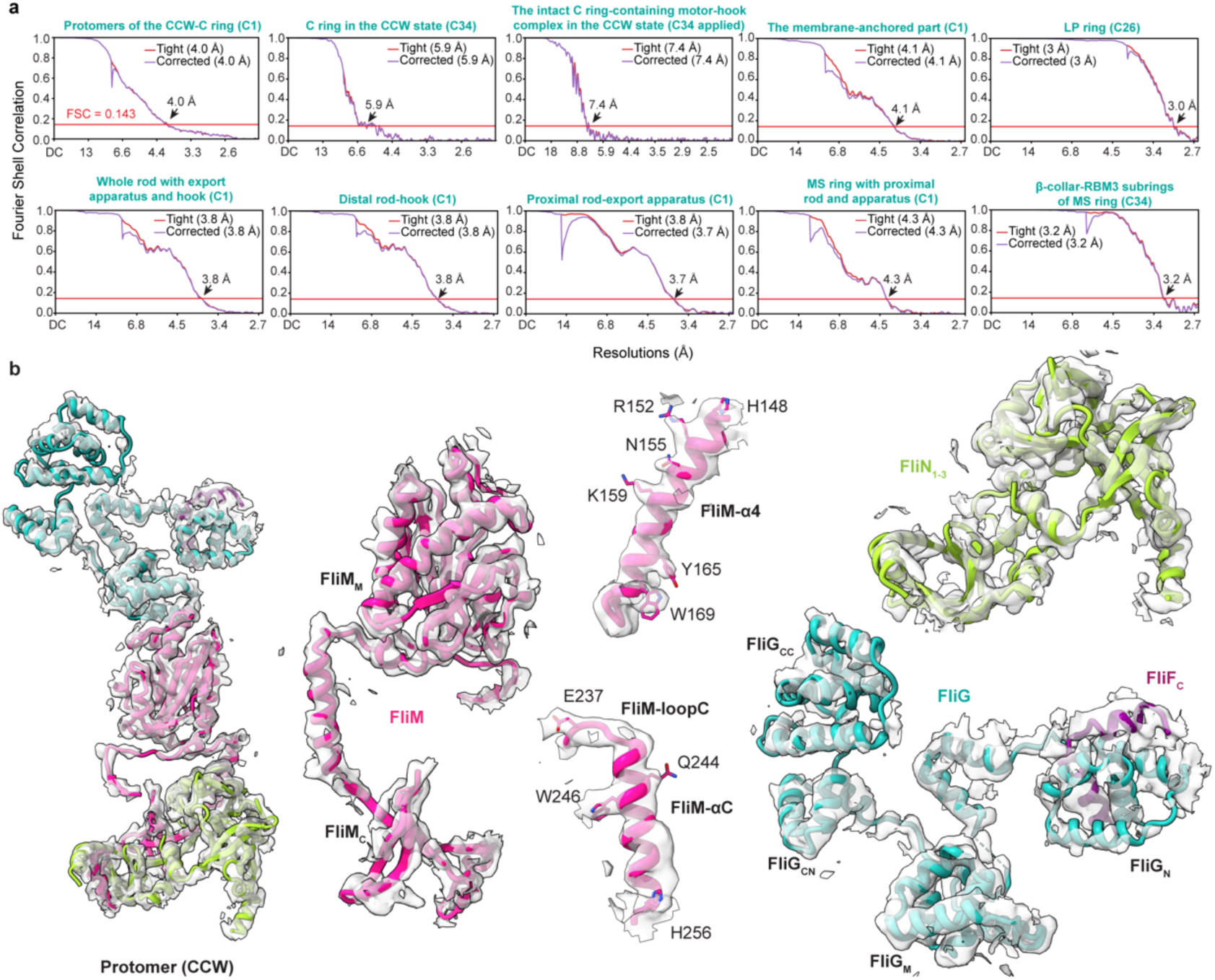
FSC curves and representative cryo-EM density maps of the components of the CCW-C ring. **a**, The Fourier Shell Correlation (FSC) curves for the reconstructions of the motor-hook complex in the CCW state. The tight and corrected FSC curves of the reconstructions are illustrated. The average resolutions are estimated based on the corrected FSC curves at the FSC=0.143 criterion. **b**, Representative cryo-EM density maps of the protomer, the FliG, FliM and FliN subunits of the CCW-C ring. Residues are shown as sticks.

**Supplementary information, Figure S3.**
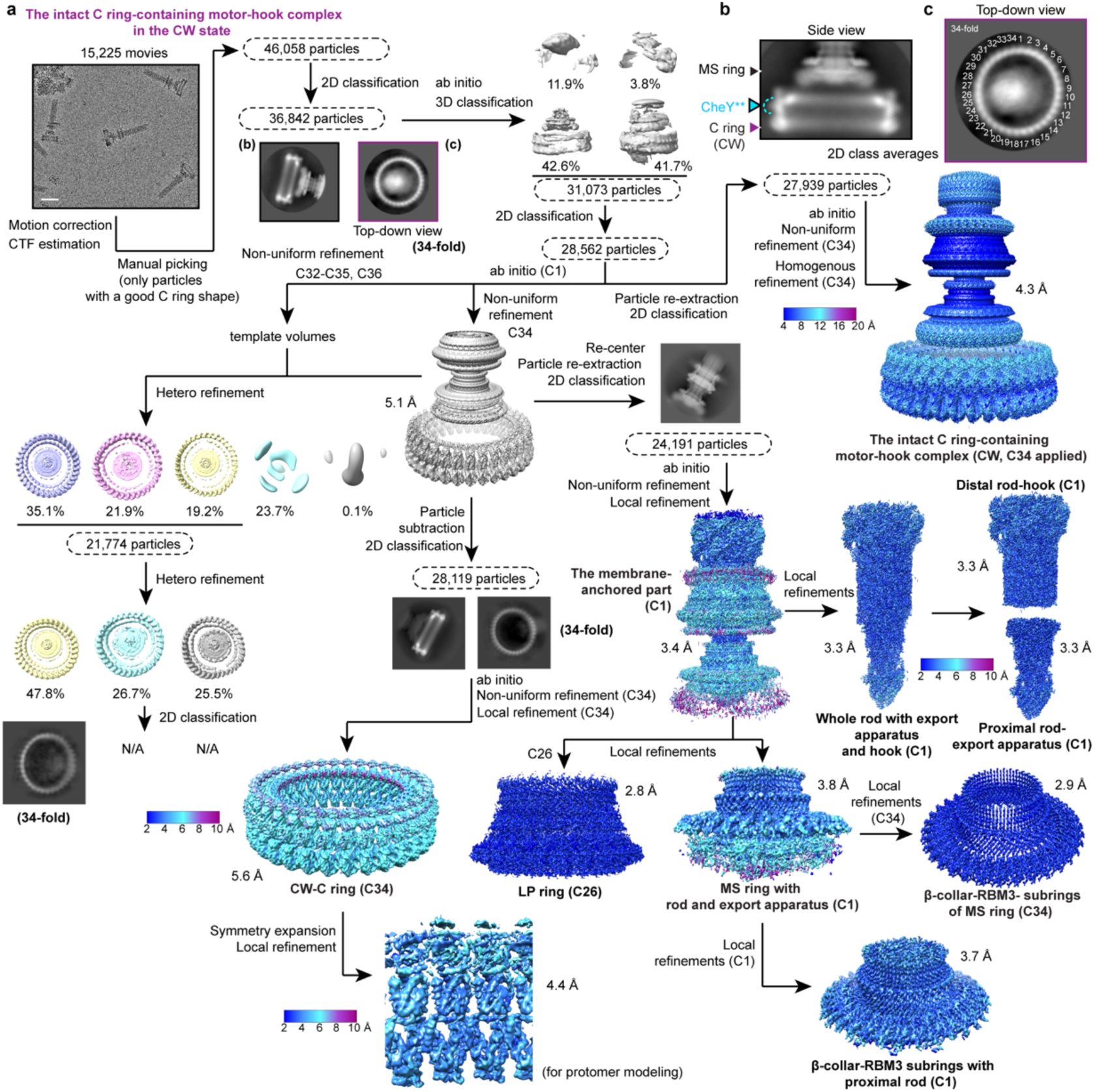
Cryo-EM data processing of the flagellar motor-hook complex in the CW state. **a**, The flow chart for the cryo-EM data processing of the C ring-containing motor-hook complex in the CW state. The local resolution maps were calculated in cryoSPARC using two independent half maps of each reconstruction as input. All density maps were prepared using *Chimera* and *ChimeraX*. Scale bar for the cryo-EM micrographs, 50 nm. N/A, not available. **b**, A representative 2D class average of the C ring in the CW state with the MS ring from the side view. **c**, A representative 2D class average of the CW-C ring from the top-down view.

**Supplementary information, Figure S4.**
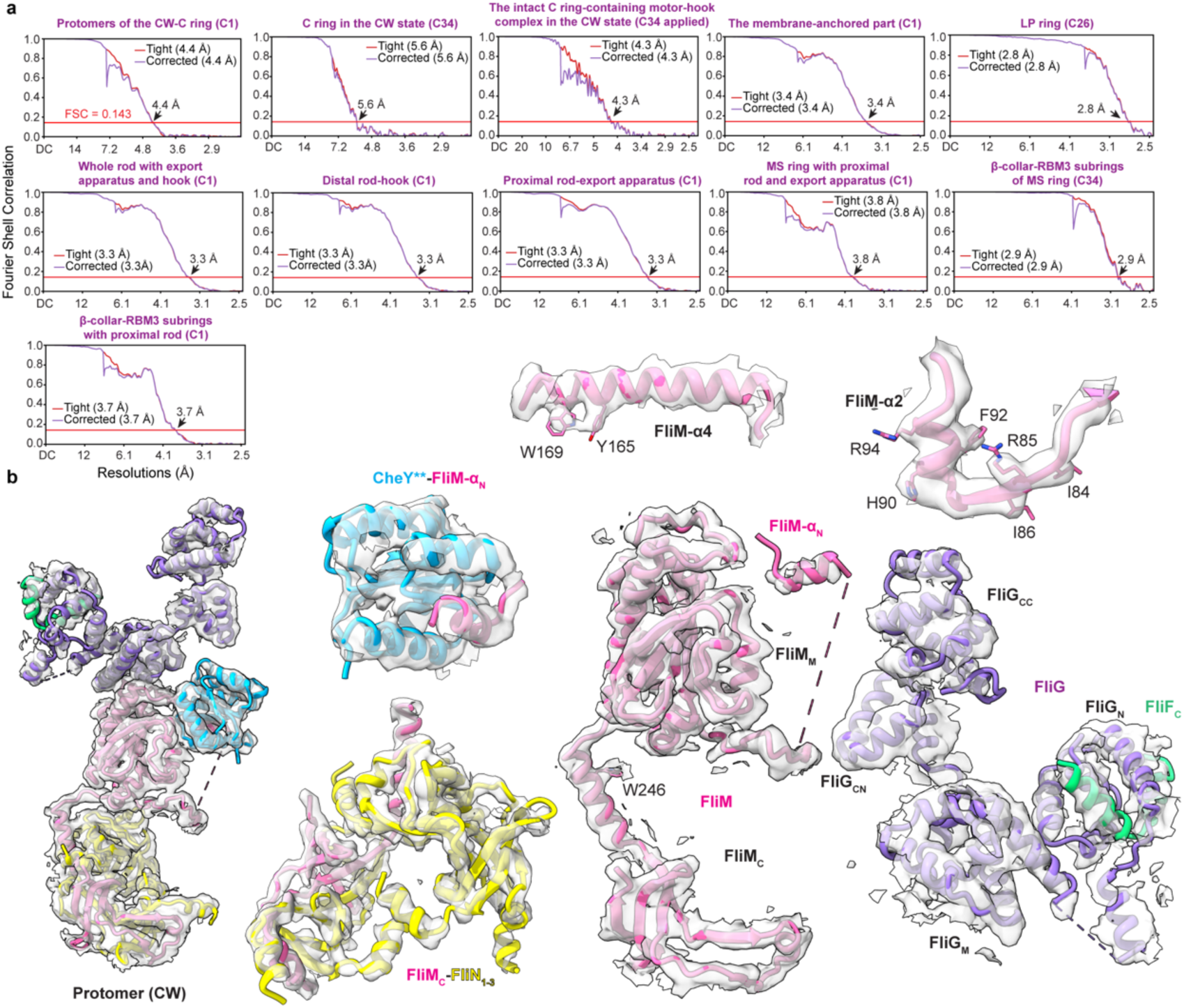
FSC curves and representative cryo-EM density maps of the components of the CW-C ring. **a**, The FSC curves for the reconstructions of the motor-hook complex in the CW state. The tight and corrected FSC curves of the reconstructions are illustrated. The average resolutions are estimated based on the corrected FSC curves at the FSC=0.143 criterion. **b**, Representative cryo-EM density maps of the protomer, the FliG, FliM and FliN subunits of the CW-C ring. Residues are shown as sticks.

**Supplementary information, Figure S5.**
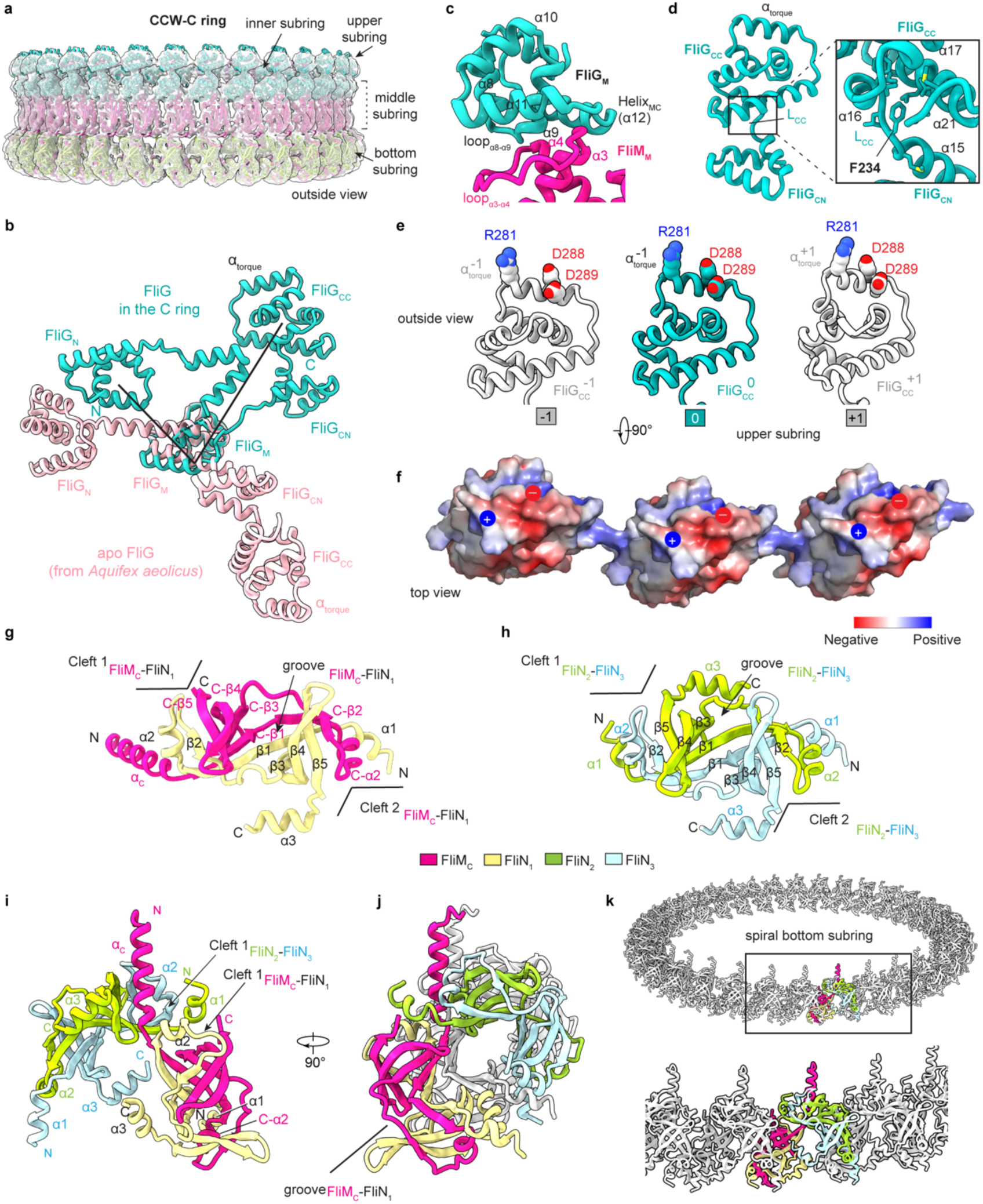
The structure and inter-subunit interactions of the CCW-C ring. **a**, The locally refined cryo-EM density map of the CCW-C ring with C34 symmetry. The four subrings of the CCW-C ring are labeled as indicated. The structural model of the CCW-C ring is colored as in Fig 2a. **b**, Structural superimposition of the structure of FliG in the CCW-C ring with the apo structure of FliG from *A*. *aeolicus*^19^ (PDB ID: 3HJL) through the FliG_M_ domains. **c**, Detailed interactions of FliG_M_ with FliM_M_ in the protomer. **d**, The detailed interactions of F234 with the FliG_CC_ domain in the CCW-C ring. **e-f**, Side view (**e**) of the conformation of the α_torque_ helices and top view (**f**) of the surface electrostatic potential of the upper subring in the CCW-C ring. The positively charged residue R281 and negatively charged residues D288 and D289 are shown as sticks and labeled as indicated (**e**). The surface is colored in blue indicates the positively charged residues and red indicates the negatively charged residues in (**f**). **g-h**, Structures of the FliM_C_-FliN_1_ heterodimer (**g**) and the FliN_2_-FliN_3_ homodimer (**h**). The FliM_C_, FliN_1_, FliN_2_ and FliN_3_ subunits are colored in red, wheat, green and light blue, respectively. **i**, Inside view of the structure of the FliM_C_-FliN_1-3_ tetramer in the CCW-C ring. **j-k**, Cross-section (**j**) and side (**k**) views of the spiral structure for the bottom subring of the CCW-C ring.

**Supplementary information, Figure S6.**
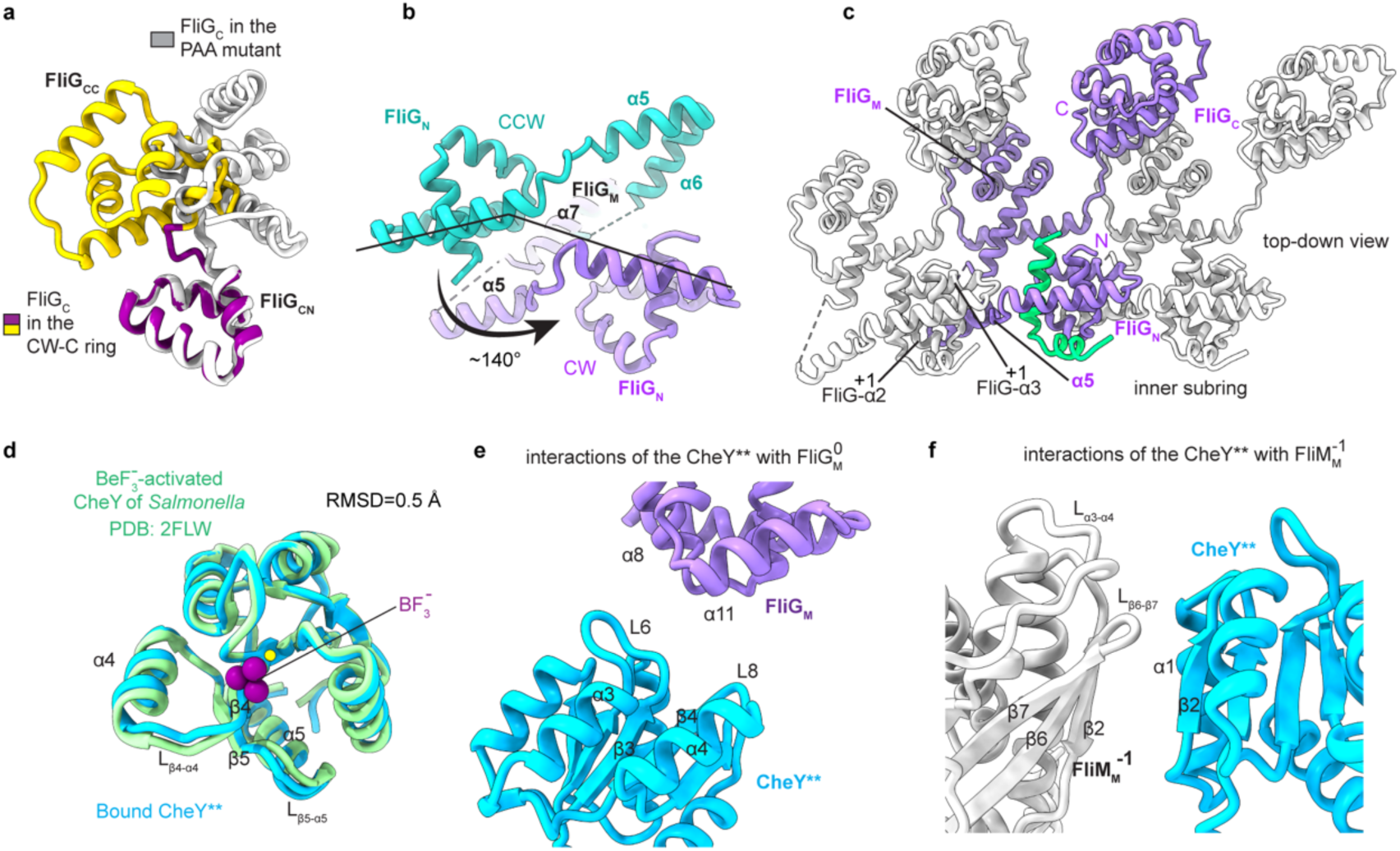
The inter-subunit interactions of the CW-C ring. **a**, Structural comparison of the FliG_C_ domains in the CW-C ring and in the crystal structure of the FliG PAA mutant (PDB ID: 3AJC)^22^. The FliG_CN_ and FliG_CC_ domains in the FliG structure from the CW-C ring are colored in purple and yellow, respectively. The FliG_C_ domain in the crystal structure is colored in grey. **b**, Structural comparison of the FliG subunits in the CCW- and CW-C rings through structural superimposition via the FliG_M_ domain. The conformational change of the FliG_N_ domain is indicated by a black arrow. **c**, Inter-subunit interactions of the FliG subunits in the CW-C ring. The FliG_N_ domain extends its α5 helix to interact with the α2 and α3 helices of the FliG^-1^ subunit of the next protomer. **d**, Structural comparison of the C ring-bound CheY** with the BF^-^ -activated CheY from *S*. Typhimurium. The structural models of CheY** and the BF^-^ -activated CheY (PDB ID: 2FLW)^35^ are colored in blue and green, respectively. The ligand BF^-^ is represented as purple spheres. **e**, Detailed interactions of CheY** with FliG_M_. **f**, Detailed interactions of CheY** with the FliM^-1^ subunit.

**Supplementary information, Figure S7.**
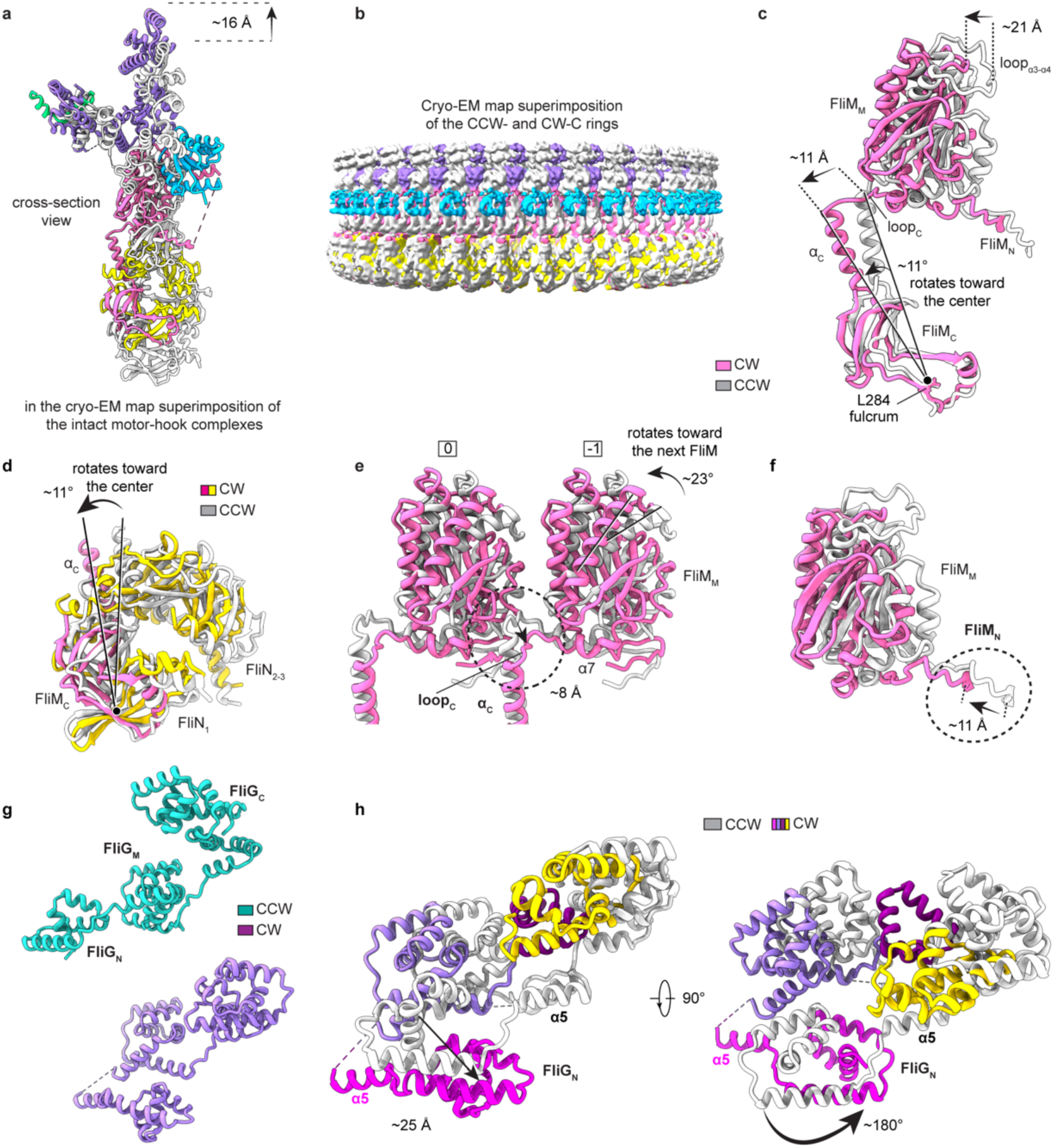
The CheY**-induced conformational changes of the C ring. **a**, Cross-section view of the structural comparison of the protomers of the CCW- and CW-C rings in the cryo-EM map superimposition of the motor-hook complexes. The protomers in the CCW- and CW-C rings are colored in grey and as in Fig 4a, respectively. **b**, Side view of superimposition of the locally refined density maps of the CCW- and CW-C rings. The density map of the CCW-C ring is colored in grey. The density map of the CW-C ring is colored according to the CW-C ring model shown in Fig 4. **c**, The conformational changes of FliM in the structural superimposition of the CCW-and CW-C rings. **d-e**, The conformational changes of FliN subunits (**d**) and the loop_C_ of FliM_C_ (**e**) in the structural superimposition of the CCW- and CW-C rings. The FliN subunits from the CW-C ring are colored in yellow (**d**). **f**, The conformational changes of FliM_N_ domains. Conformational changes are indicated by black arrows and a dashed circle (**c-f**). The FliM subunits from the CCW- and CW-C rings are colored in grey and pink, respectively (**c-f**). **g**, Structural comparison of the overall structures of FliG in the CCW- and CW-C rings. FliG in the CW-C ring (medium purple) has a more compacted conformation than that in the CCW-C ring (cyan). **h**, Inside (left) and top (right) views of the domain rearrangement of FliG in the CCW- and CW-C rings. The FliG_N_ domain rotates ∼180° towards the prior protomer and moves downward by ∼25 Å.

**Supplementary information, Figure S8.**
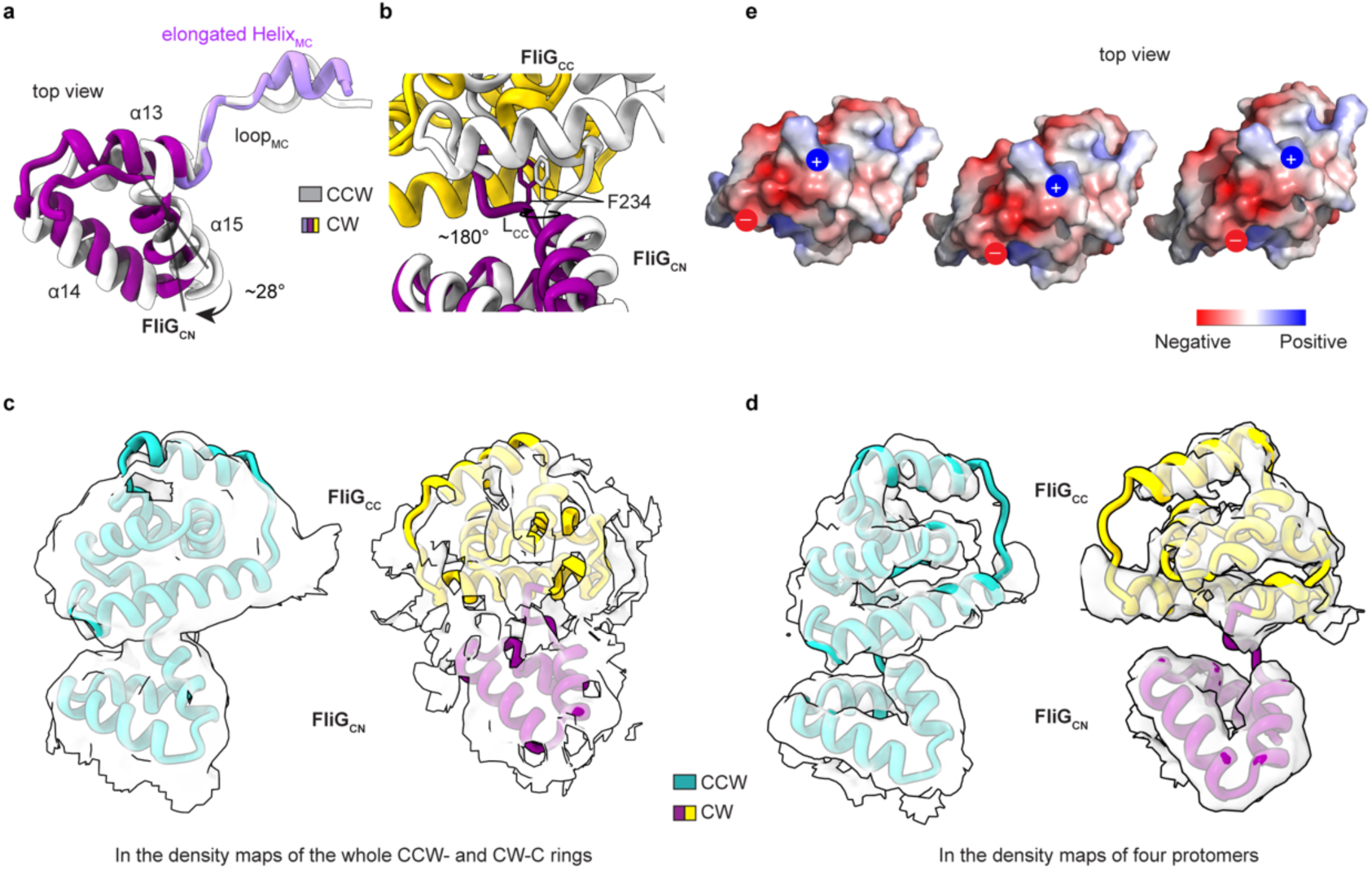
The conformational changes and cryo-EM density maps of the FliG_C_ domain. **a-b**, Enlarged views of the CheY**-induced rotation of FliG_CN_ on the surface of FliG_M_^-1^ (**a**) and the conformational changes of the hinge loop L_CC_ (**b**) in the structural comparison in Fig 5f. **c-d**, Densities of the FliG_CN_ and FliG_CC_ domains of the CCW- and CW-C rings in the density maps of the C rings (**c**) and in the locally refined maps of four protomers (**d**). **e**, The surface electrostatic potential of the upper subring in the CW-C ring. The surface is colored in blue indicates the positively charged residues and red indicates the negatively charged residues.

**Supplementary information, Figure S9.**
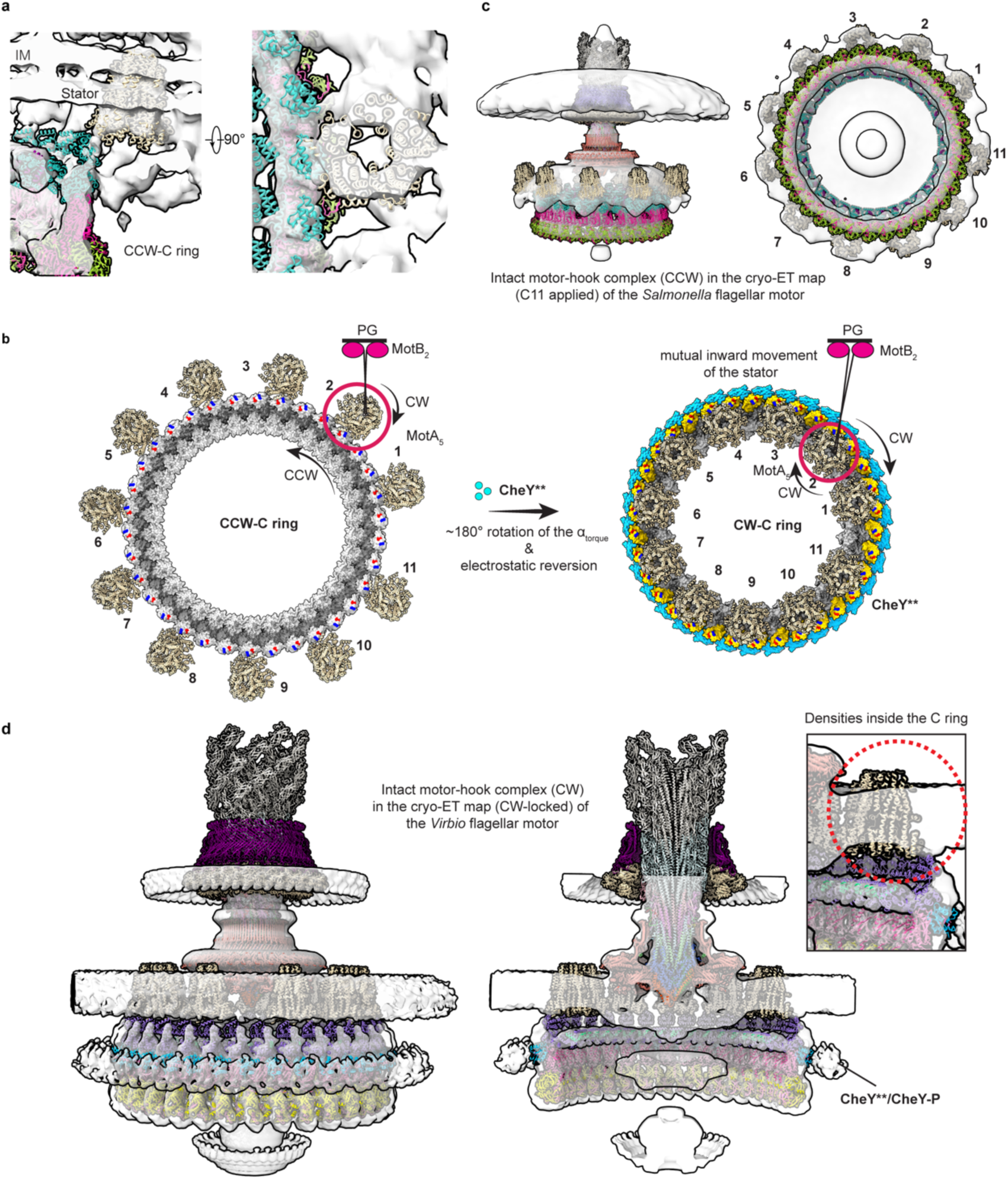
The structural analyses of the relocation process of the stators for rotational switching. **a**, Cross-section (left) and top (right) views of structural modeling of the CCW-C ring and the MotA_5_ pentamer of a stator of *S*. Typhimurium into the cryo-ET density map of the flagellar motor of *Helicobacter pylori* (EMD-25123) in the default CCW state^36^. The structural models of the CCW-C ring and the MotA_5_ pentamer are colored as in Fig 2a and wheat, respectively. **b**, Structural models of the interactions of the C ring with 11stator units in the intact *Salmonella* motors in the CCW (left) and CW (right) states. **c**, The pseudo-atomic model of the intact motor-hook complex containing CCW-C ring and 11 stators in the cryo-ET density map (EMDB: EMD-3154) of the flagellar motor of *S*. Typhimurium^39^. **d**, The protruding densities in the inner membrane, which are above the inner side of the upper subring of the CW-C ring, for accommodation of the stator units in the cryo-ET density map (EMDB: EMD-21837) of the flagellar motor of *V. alginolyticus* in the CW state^27^. The pseudo-atomic model of the intact motor-hook complex containing CW-C ring and 11 stator units, which was obtained in Fig 6d, was fitted into the cryo-ET density map, and colored as indicated. The protruding densities were ignored and not annotated in the previous study.

**Supplementary information, Figure S10.**
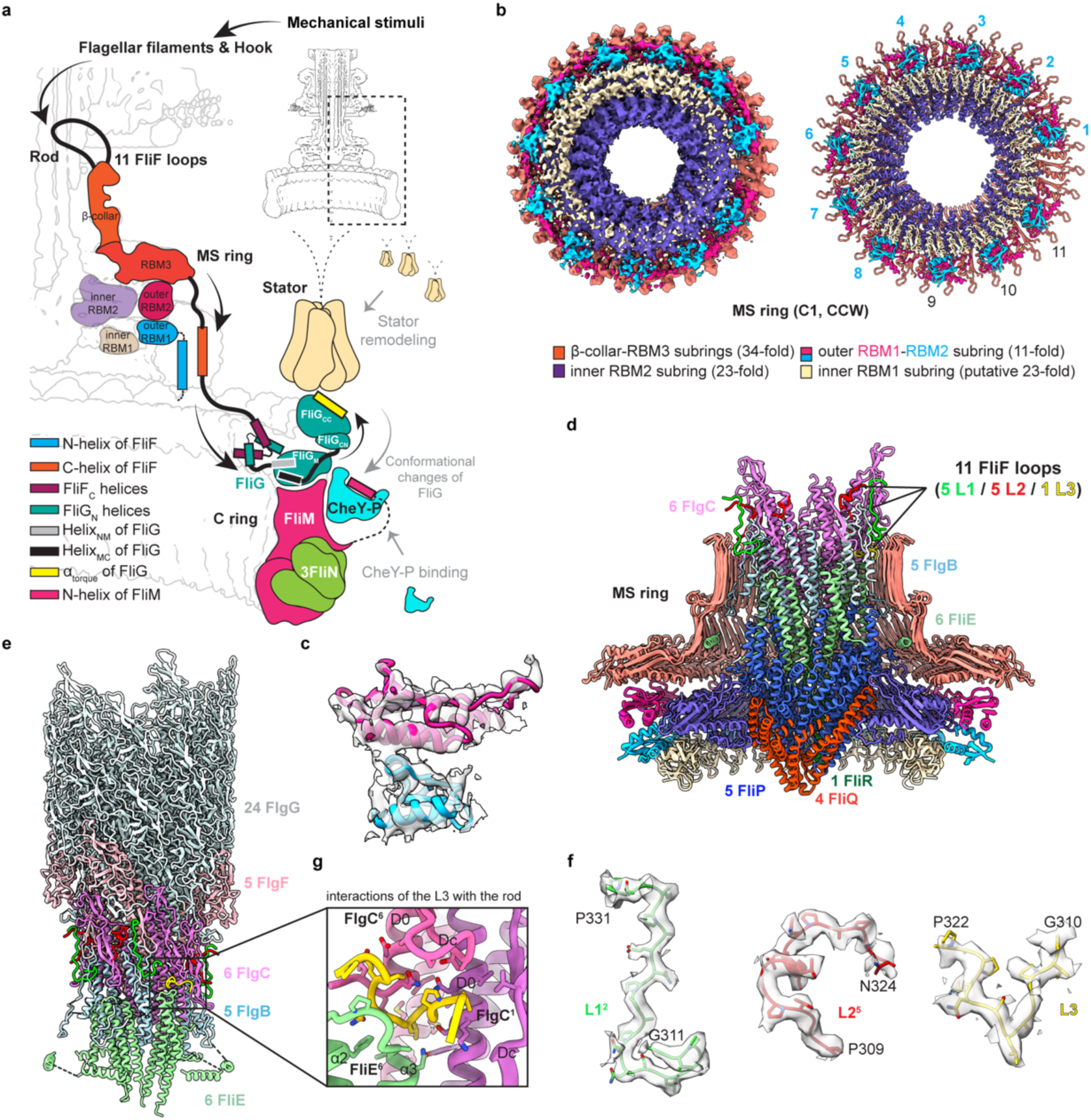
Schematic diagram for potential mechanosensing mechanism of the flagellar motor, and interactions of the MS ring with the rod in the C ring-containing motors. **a,** Schematic diagram for the role of the C ring in the mechanosensing of the bacterial flagellar motor. The environmental mechanistic stimuli likely lead to the conformational changes of the filaments, hook and rod. The 11 FliF peptide loops that are bound on the rod surface induce the conformational changes of the MS ring. the C ring adjusts the conformations of the FliG_CC_ domains via the Helix_NM_-Helix_MC_-FliG_CN_-FliG_CC_ module cascade upon the conformational changes of the MS ring to regulate the loading of the stators, as well as the recruitment of the CheY-P towards the C ring. **b**, Bottom views of the cryo-EM density map (left) and structure (right) of the MS ring from the motor-hook complexes in the CCW state. The 4.3-Å density maps of the MS ring were reconstructed with C1 symmetry as shown in Fig **S1a** is illustrated. The 34-fold β-collar-RBM3, 23-fold inner RBM2, 11-fold outer RBM2, 11-fold outer RBM1 and putative 23-fold inner RBM1 subrings are colored in salmon, purple, red, blue and wheat, respectively. In this model, the putative 23-fold RBM1 subring and 3 of the 11 outer RBM1-RBM2 domains are modelled (labelled with black numbers) according to the density map but are not built in the final structure. **c**, Representative density maps of the outer RBM1 and RBM2 domains in the CCW-C ring. **d**, Cross-section view of the MS ring with the embraced proximal rod and the export apparatus in the motor-hook complex in the CCW state. The L1, L2 and L3 loops of FliF, which are extended from the MS ring, are highlighted in green, red and yellow, respectively. **e**, Interactions of the 11 FliF peptide loops from the MS ring with the rod. The subunits of the proximal rod are colored and labeled as indicated. All residues (P309-N324) of L2 were precisely modeled in the C ring-containing motor-hook complexes. The 11 loops are packed in the same right-handed helical manner as the rod subunits. **f**, Representative density maps of the L1, L2 and L3 loops. The L3 loop is observed in both the motor-hook complexes in the CCW and CW states. The models and density maps of the L3 loop from the CW-biased motor are illustrated. **g**, Detailed interactions of the L3 loop with the rod subunits. The interacting residues are shown as sticks.

## Notes

### Competing Interest Statement

The authors have declared no competing interest.

